# Metabolic modeling predicts synergistic growth benefits between arbuscular mycorrhizal fungi and theoretical N_2_-fixing rhizobia symbiosis in maize

**DOI:** 10.1101/2025.01.28.635303

**Authors:** Joshua A.M. Kaste, Rourou Ji, Patrick Sydow, Ruairidh J. H. Sawers, Megan L. Matthews

**Affiliations:** Department of Civil and Environmental Engineering, Grainger College of Engineering, University of Illinois Urbana-Champaign, 205 N Mathews Ave, Urbana, IL 61801; Department of Plant Science, College of Agricultural Sciences, The Pennsylvania State University, 102 Tyson Building, University Park, PA 16802; Carl R. Woese Institute for Genomic Biology, University of Illinois Urbana-Champaign, Urbana, IL 61801

**Keywords:** Flux Balance Analysis, Arbuscular Mycorrhizal Fungi, N2 fixation

## Abstract

Engineering a novel N_2_-fixing rhizobia symbiosis in cereal crops is a strategy being pursued to improve agricultural sustainability. However, if such a symbiosis were introduced, it would have to be economically viable in the context of existing nutrient acquisition strategies, including the existing symbiosis with arbuscular mycorrhizal fungi (AMF) that the vast majority of plants already engage in. This raises the question of how the metabolic costs and benefits from these separate symbioses that have partially overlapping functions might impact nutrient status and subsequent plant growth. To address this, we developed metabolic models describing how the relative growth rate of *Zea mays* is impacted by the AMF *Rhizophagus irregularis* and a hypothetical N_2_-fixing symbiosis with *Bradyrhizobium diazoefficiens* both in isolation and in tandem. To validate the AMF component of our model, we conducted field evaluation of mutant AMF-incompatible maize hybrids and found that the empirically measured AMF-mediated growth benefit agreed well with our model’s predictions. Our model of the rhizobium symbiosis predicted that the lower N content of cereal crops makes the relative growth rate cost associated with acquiring nitrogen from N_2_-fixing rhizobia smaller than in legumes. Finally, our model also predicted positive synergies between rhizobia and AMF under nutrient-limited conditions but negative synergies under nutrient, particularly phosphorus, replete conditions. These findings indicate that these bioengineering strategies could improve cereal crop yields and may achieve greater gains in tandem, but soil nutrient status of target sites as well as the nitrogen requirements of specific varieties should be considered.

## Introduction

Agricultural production poses a huge challenge to global efforts to address the intertwined problems of anthropogenic climate change and sustainable agricultural development. Approximately 19% of greenhouse gas emissions can be attributed to agriculture and deforestation (1). CO_2_ emissions associated with nitrogen fertilizer production and the release of N_2_O from agricultural fields post-application contribute heavily to these emissions (2–4). Additionally, these fertilizers represent a substantial portion of the cost of agriculture and recent volatility in fertilizer prices has exacerbated food insecurity in Africa (5). Increases in the cost of urea in Africa specifically have been linked with increased deforestation due to expanded agricultural land use (6). For these various reasons, researchers are looking to develop alternatives to synthetic fertilizers to allow cultivators, particularly in developing economies, to achieve high crop yields sustainably and profitably.

One promising approach to tackle these challenges is to engineer associations with N_2_-fixing bacteria into crops that do not currently have this capacity (4,7,8). If accomplished, this would allow non-leguminous plants, in symbiosis with N_2_-fixing rhizobia, to fix nitrogen instead of relying on external nitrogen application. This approach could be most effective in a cereal crop with low nitrogen content, like maize (*Zea mays*), since the costs incurred in this symbiosis by the plant host – e.g., the provisioning of dicarboxylates to the rhizobia partner to fuel nitrogen fixation – increase with the plant’s nitrogen requirements. Due to this lower nitrogen requirement, a previous analysis predicted that, compared with a legume like *Glycine max*, maize engineered to host rhizobia may provide less resources to this symbiosis when similar levels of N are available, therefore reducing the negative effect on relative growth rate (RGR) (9). However, this reduced RGR penalty was inferred through a simple comparison of the nitrogen content of maize and soybean, rather than through a full metabolic modeling analysis. This analysis also did not account for changes in tissue carbon and nitrogen content during maize development, which may result in the growth penalty varying over time.

In both legumes and non-leguminous crops, these real and potential symbioses with rhizobia exist, or would exist, alongside symbioses with arbuscular mycorrhizal fungi (AMF) (10). The plant-AMF symbiosis can confer a variety of benefits to plants, including increased access to soil nitrogen and phosphorus through symbiotic exchange with AMF, which can reduce the amount of fertilizer necessary for optimal crop growth (11). Because of this, enhancement of this existing capacity for crops to associate with and take up nutrients from AMF has also become a subject of intense research (12). There is reason to believe that the plant-rhizobia and plant-AMF symbioses may complement each other. The potential growth improvements from N_2_-fixation by rhizobia in cereal crops may be limited in some soils by low phosphorus availability. AMF improve plant phosphorus uptake (13) and could allow plants to fully realize rhizobia-mediated growth benefits in phosphorus-poor soils. A meta-analysis on the effects of AMF-inoculation on plant growth found that in laboratory studies of AMF fungi, N_2_-fixing forbs benefited more from AMF-inoculation than C4 grasses did, suggesting possible synergy between these symbiotic associations (14). Positive synergy between rhizobia and AMF has also been reported in white clover (15), and another study found that rhizobia also affected the expression of several mycorrhizal genes, including those involved in nutrient transfer to host plants, indicating that partner species can also impact each other’s molecular phenotypes (16). Collectively, these data illustrate the diverse molecular mechanisms and transcriptional responses associated with the synergistic benefits of multiple mutualists. These observations make an analysis of this three-species symbiosis and its effects on plant growth highly relevant.

To perform this analysis, models describing (i) the hypothetical symbiosis between a cereal crop and a rhizobium, and (ii) the symbiosis between a cereal crop and AMF first need to be constructed, before combining them to analyze synergies and antagonisms in the full system. Plant-rhizobia metabolic models have been developed describing legumes, but have not explored the costs and benefits of potential cereal symbioses (9) and plant-AMF models have described individual components of the symbiosis (17–19), but have stopped short of explicitly connecting plant and fungal metabolism via genome-scale models. In maize specifically, modeling has been done that uses transcriptomic data to predict metabolic alterations in the plant in response to AMF-inoculation, but this is done only with a maize model and without explicitly modeling metabolite exchanges between the plant and AMF (20,21). To quantify the metabolic costs and benefits of these proposed strategies, a modeling study explicitly describing the metabolite exchanges between a cereal crop, AMF symbionts, and potential N-fixing rhizobia symbionts both in isolation and in tandem is needed.

In this study, we use genome-scale models of the metabolic networks of the C4 grass *Zea mays*, the N_2_-fixing rhizobium *Bradyrhizobium diazoefficiens,* and the AMF *Rhizophagus irregularis*, to characterize the costs and benefits of rhizobia and AMF on crop growth in isolation and together. First, we develop a model of maize central metabolism, adapted from a model of *Arabidopsis thaliana* core metabolism (22) that accurately reproduces essential growth characteristics of the plant. We then connect this with a genome-scale model of *R. irregularis* (23) to simulate the exchange of nutrients between maize and AMF. We generate and present experimental field data comparing hybrid *Zea mays* genotypes that are capable and incapable of associating with AMF and show that our model predictions align with our empirical observations when accounting for available soil nitrogen and phosphorus. Next, we combine our *Zea* mays model with a model of the N_2_-fixing rhizobium *Bradyrhizobium diazoefficiens* (24) and use it to calculate the carbon cost of biological nitrogen fixation in this system. Our results confirm previous predictions that the carbon cost should be lower than it is in soybean, but we also find that this cost would vary across plant development. Using these estimated carbon costs, we perform a cost-benefit analysis that shows that the incorporation of the N_2_-fixing rhizobia into the maize system would be more cost-effective than synthetic N fertilizer applications. Finally, by combining the models describing the AMF and rhizobia associations, we predict strong synergy between these strategies under low phosphorus and nitrogen conditions in terms of their positive impact on plant RGR (see **Figure S1** for a graphical representation of the model). Under phosphorus-replete soil conditions, however, we predict antagonism between these approaches.

## Results

### Construction and validation of a multi-tissue model of maize central metabolism

For this study, a reconstruction of *Zea mays* metabolism is needed that 1) represents the separation of the mesophyll and bundle sheath cells to accurately represent C4-associated metabolic costs and 2) is computationally tractable in terms of model size. While several generic C4 and *Zea mays* specific reconstructions exist (25–28), none of them address both of these needs. We constructed a multi-tissue metabolic model of maize central metabolism using a bottom-up reconstruction approach using the central metabolism of *Arabidopsis thaliana* as a base (22). The model features a biomass equation representative of maize (27), separation of the mesophyll and bundle-sheath cells to simulate C4 photosynthetic fluxes, and separate root and shoot tissue compartments connected by inter-tissue metabolite exchanges based on a previous study (20). Where necessary, additional biochemical reactions were added to the model from KEGG and other prior literature sources to allow the synthesis of all biomass components **(Dataset S1)**.

We compared key outputs of our model, focusing on measures of efficiency and characteristics of C_4_ photosynthesis, to literature estimates and known biochemistry and found them highly consistent **(Table 1)**(29–32). Taken together, these validation results indicate that the model captures key photosynthetic details and the overall energetic efficiency of maize central metabolism. Local sensitivity analysis was performed on this base plant model and the other models presented in this study and shows that outputs of interest are robust to small variations in most key parameter values, though the necessary AMF biomass parameter notably shows high sensitivity (**Figure S2)**. As we show later, however, an uncertainty analysis shows that, due to the relatively small error associated with our estimate of this parameter, it does not have an outsized influence on our results or conclusions. Our model’s bundle sheath cells also appears to rely somewhat less on cyclic electron flow (CEF) as compared with those measured in the leaves of another C_4_ plant, *Setaria viridis*, though this may be due to inter-species differences.

**Table 1:** Photosynthetic and physiological parameter predictions from the model of maize along with qualitative and quantitative data and corresponding references.

| Metric | Modeled | Measured | Reference |
| --- | --- | --- | --- |
| Organic Carbon Transport from Mesophyll to Bundle Sheath | Malate | Malate | (30) |
| Carbon Skeleton Transport from Bundle Sheath to Mesophyll | Pyruvate | Pyruvate | (30) |
| Decarboxylated Species in Bundle Sheath | Malate | Malate | (30) |
| Relative Growth Rate ($\text{g gDW}^{-1} \text{d}^{-1}$) | 0.082 | 0.04 – 0.08 | (32) |
| Quantum Yield of Photosynthesis ($\text{mol CO}_2 \text{ mol photons}^{-1}$) | 0.0541 | 0.052; 0.059 | (79,80) |
| Percent of Bundle Sheath PSI Electrons Supplied by Cyclic Electron Flow (%) | 65% | 84%<sup>a</sup> | (31) |

<sup>a</sup>Note that this measurement was made in a non-maize C<sub>4</sub> plant (*Setaria viridis*).

### Experimentally validated plant-AMF model relates biomass improvements in maize to nutrient and metabolite exchanges

To understand the relationship between the nutrient uptake benefits mediated by AMF and biomass benefits observed in plants, we built a multi-species plant-AMF model by combining our *Zea mays* metabolic model with an existing genome-scale model of the AMF *Rhizophagus irregularis* (22,23). Prior literature suggests that plants allocate carbon to AMF symbionts in the form of hexoses, primarily glucose (33,34), and lipids, most likely in the form of palmitate (35,36). In our plant-AMF model, these carbon sources are exchanged for nitrogen (represented as NH_4_ in our model) and phosphorus taken up by the AMF. The plant model can also take up nitrogen and phosphorus itself from the surrounding soil, with the maximum rate at which it can obtain these nutrients varied from 100%, representing the uptake rate needed for plant growth to no longer be limited by that nutrient’s availability, down to 0%. Key parameters for this model include the percent improvement of external nitrogen and phosphorus uptake rate that the plant-AMF system can achieve compared to the base plant uptake, and the amount of fungal biomass necessary to achieve these uptake rate improvements (**Table S1**). We inferred these parameters from literature sources (37–39), but they have moderate-to-high uncertainty associated with them. To account for the impact of this uncertainty on our model predictions, we used a Monte Carlo sampling approach.

The relative maize growth benefits from AMF-mediated nutrient uptake are predicted to be maximal in nitrogen-replete and phosphorus-depleted conditions (**Figure 1A)**. Accordingly, because our model allocates carbon investment to the fungus to maximize plant RGR, carbon investment is maximized under those same conditions. This carbon investment decreases, eventually to zero, as nitrogen and phosphorus availability increases and plant uptake alone becomes sufficient to achieve maximum, light-limited plant RGR **(Figure 1B).** These predictions are consistent with the trade-balance model of the AMF symbiosis, which describes differing levels of growth benefit conferred by AMF on plants in terms of nutrient availability, relative nutrient uptake capacities of plants and AMF, and the biomass composition of plant and fungal tissues (13). By comparing the carbon allocated by the plant to the fungus and the nutrients received in return in our model, we can estimate the stoichiometry of carbon and nutrient exchange between the plant and AMF partner. The values vary with external nutrient levels, which we show in **Dataset S2**. Under strictly P limiting conditions, the stoichiometry of carbon-phosphorus exchange is 15 g C g P^-1,^ and under strictly N limiting conditions, the stoichioetry of carbon-nitrogen exchange is 7 g C g N^-1^ (**Dataset S2).**

**Figure 1:**
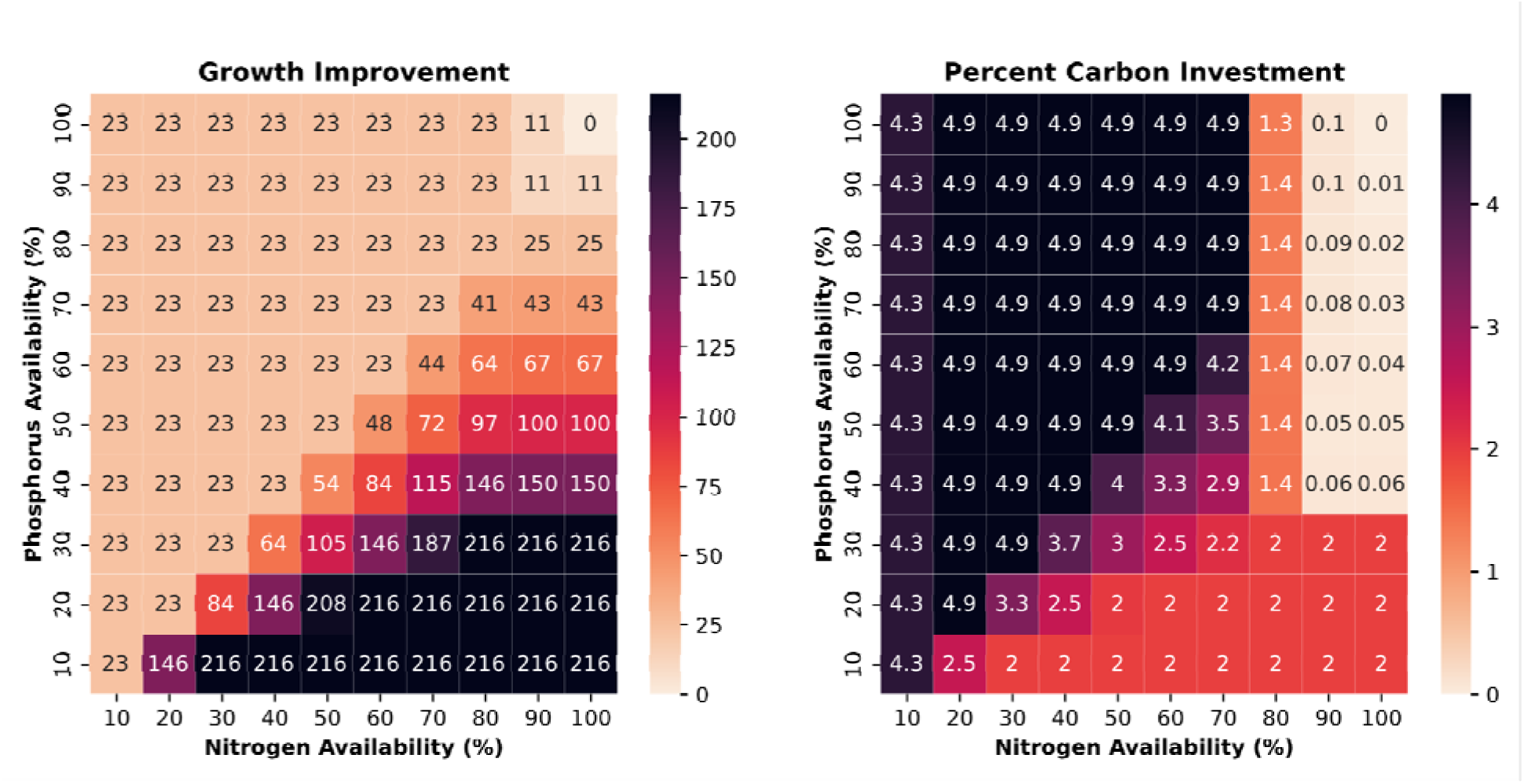
Average growth improvement and carbon investment predictions when comparing with– and without-AMF models when maize biomass is representative of seedling C:N ratios. **(A)** Percentage growth improvement of with-AMF vs. without-AMF models of maize as a function of phosphorus and nitrogen availability. **(B)** Predicted carbon investment in AMF as a percentage of net CO_2_ assimilation in the plant as a function of phosphorus and nitrogen availability. For plots of growth improvement and carbon investment when the plant biomass is representative of plants at the jointing or silking developmental stages, refer to **Figures S4-5**.

Our local sensitivity analysis **(Figure S2B-C)** shows that the stoichiometry predictions from our model vary linearly with the assumed amount of AMF dry weight needed per unit plant dry weight. This means that the accuracy of our estimate of the AMF dry weight needed will determine how reasonable the g C g N^-1^ and g C g P^-1^ estimates are. Using data on the hyphal length density of AMF under conditions where phosphorus uptake benefits are observed in plants (37) and literature estimates on the density and fresh-to-dry weight ratio of fungal tissue (39), we estimated that AMF biomass amounting to 9% of the total plant biomass is necessary to support maximum nutrient uptake benefits. This is slightly higher but qualitatively similar to a previous study’s estimate of ∼6% (35). With this assumption, the range of carbon investment predicted by the model, 0 to 5% of plant carbon (**Figure 1B**), is consistent with values reported in the literature (2.3 to 10% for *R. irregularis* (40)).

To assess whether the growth-rate improvements predicted by our model are consistent with those observed in the field, we used empirical time-series measurements taken from field-grown maize hybrids with and without a mutation in the gene *Castor* that results in an inability to associate with AMF (41). These plants were grown in phosphorus-replete (100%+ of necessary P) and nitrogen-limiting (∼84% of necessary N) soil conditions (**Dataset S3**). The 95% confidence interval (CI) on the observed RGR benefit when comparing AMF– and AMF+ plants in this experimental dataset is 11 – 47%, which compares well with the predicted range of 15 – 32%. **(Figure 1A, Dataset S2)**. However, our model does not account for dynamic changes in biomass composition as a function of nutrient availability and symbiont presence/absence. Although the proportion of tissue biomass made up of N is not statistically significantly different between AMF+ and AMF-plants in the field data, the AMF+ plants have a ∼180% increased P content relative to the AMF-plants **(Dataset S3)**. Because of this, the model underestimates the cost, in terms of necessary P, to build a unit of biomass in the AMF+ plants. Accounting for this increased P cost results in an effective RGR benefit prediction of 8 – 18%, which overlaps with the lower range of the observed benefit. This shows that our model accurately predicts AMF-mediated growth benefits under phosphorus-replete, nitrogen-limited conditions. Because our field was N-limited, but not P-limited, to assess the accuracy of our model in P-limited conditions, we compared model outputs to a previous greenhouse experiment conducted under P-limited conditions (37). Under phosphorus-limited conditions, our model predicted a 147 – 287% improvement in RGR **(Figure 1A)**, as compared with the range of 66 – 150% improvement, depending on cultivar, in RGR in AMF+ vs. AMF-maize plants measured in a previous greenhouse study (37). If we account for this study’s observed 44% increase in biomass phosphorus concentration (37), we would instead predict an 82 – 160% improvement, consistent with what was measured. Note that (37) did not report time-series dry weight data, which is necessary to calculate RGR, so we estimated the RGR based on the final dry weight measured after 8 weeks (36) compared to a typical kernel dry weight of 2.5g.

### The predicted RGR penalty incurred by N_2_-fixation in maize is lower than in non-cereal crops and decreases across vegetative development

A model of *Bradyrhizobium diazoefficiens* (24) was connected to the maize model, with a new nodule tissue compartment, to investigate the potential benefits and drawbacks of creating an association between maize and an N_2_-fixing rhizobium. By comparing simulation results with and without the rhizobia and related plant tissues present, we quantified the RGR penalties of using N_2_-fixation in lieu of soil nitrogen uptake across a range of soil nitrogen availabilities and across several different vegetative developmental stages in maize **(Table 2**; **Figure 2)**.

**Figure 2:**
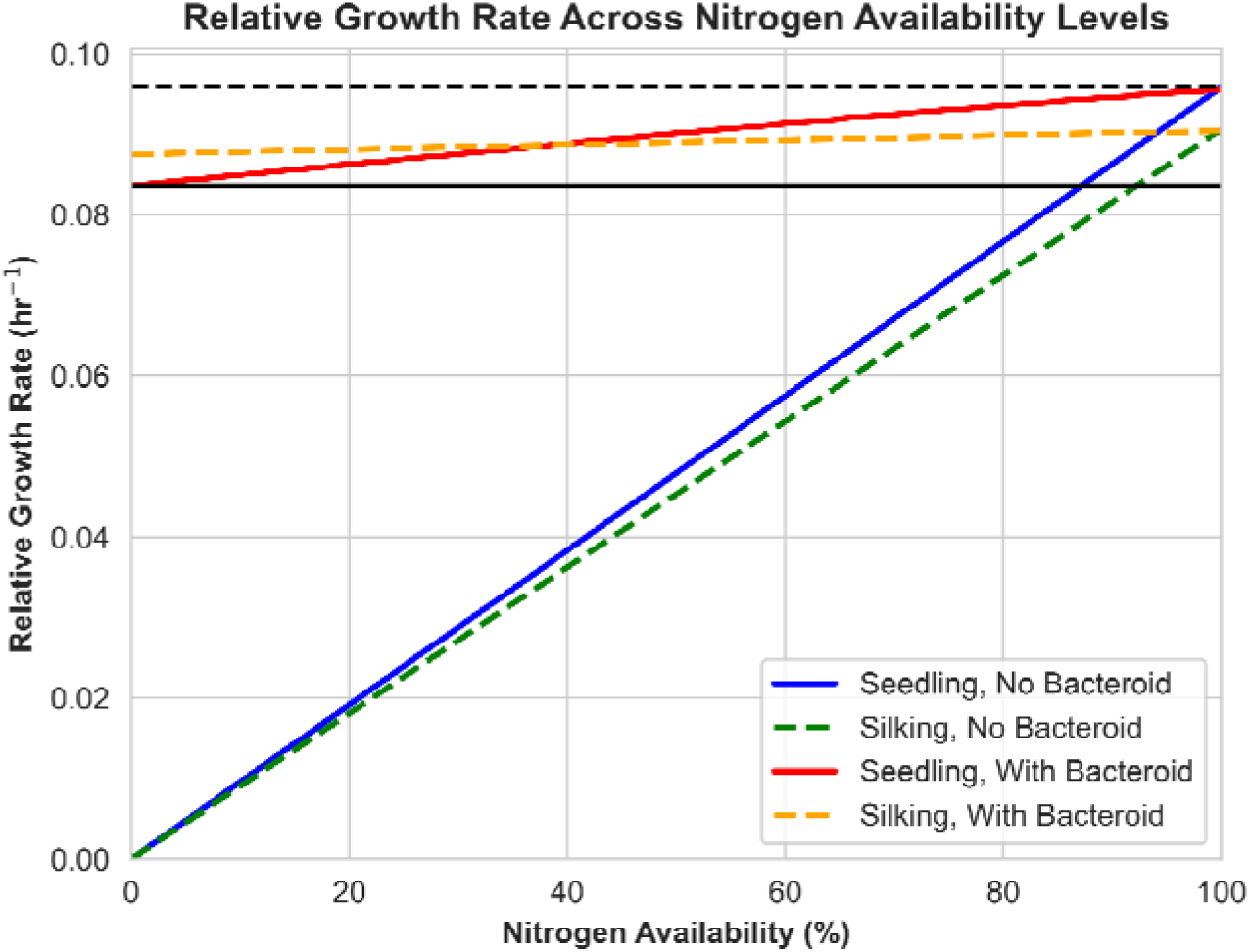
Relative Growth Rate of *Z. mays* across a range of nitrogen availability levels for models with biomass representative of maize seedlings and maize plants at silking. Solid lines represent growth predictions when using the seedling biomass equation and dashed lines represent growth when using the silking biomass equation. The dashed black line represents the maximum growth rate of the non-nodulated model when it receives all its nitrogen from the soil. The solid black line represents the maximum growth rate of the nodulated model when it fully relies on N_2_-fixation. For the relative growth rates of the models with biomass representative of the C:N ratios of maize plants at the jointing stage, see **Figure S6**.

**Table 2:**
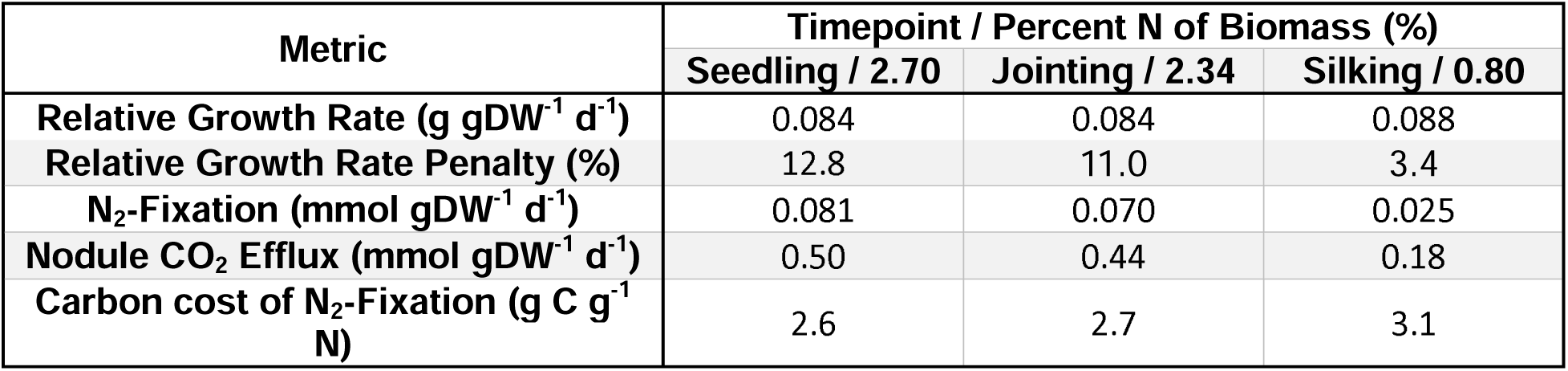
Physiological and metabolic predictions from the *Z. mays* model associated with the N_2_ fixing rhizobium *B. diazoefficiens*.

| Metric | Timepoint / Percent N of Biomass (%) |  |  |
| --- | --- | --- | --- |
|  | Seedling / 2.70 | Jointing / 2.34 | Silking / 0.80 |
| Relative Growth Rate (g gDW<sup>-1</sup> d<sup>-1</sup>) | 0.084 | 0.084 | 0.088 |
| Relative Growth Rate Penalty (%) | 12.8 | 11.0 | 3.4 |
| N<sub>2</sub>-Fixation (mmol gDW<sup>-1</sup> d<sup>-1</sup>) | 0.081 | 0.070 | 0.025 |
| Nodule CO<sub>2</sub> Efflux (mmol gDW<sup>-1</sup> d<sup>-1</sup>) | 0.50 | 0.44 | 0.18 |
| Carbon cost of N<sub>2</sub>-Fixation (g C g<sup>-1</sup> N) | 2.6 | 2.7 | 3.1 |

The RGR penalty associated with maize acquiring all of its nitrogen from N_2_-fixation versus soil uptake ranges from ∼12.8% in the seedling stage down to 3.4% at the silking stage, which is qualitatively consistent with prior estimates (9). The predicted carbon cost of N_2_-fixation ranges from 2.6 g C g N^-1^ at the seedling stage to 3.1 g C g N^-1^ at the silking stage. This decreased efficiency results from fixed non-growth associated maintenance in the nodule and bacteroid tissues with decreased use of N_2_-fixation during the silking stage when nitrogen requirements begin to decrease (**Table 2**). These values are consistent with the range of empirically measured g C g N^-1^ values in the literature for soybean (2.5 to 7 g C g N^-1^)(42).

### Value Cost Ratio comparisons indicate that a nitrogen-fixing maize crop could be profitable in both the United States and sub-Saharan Africa

To assess the economic viability of inoculating a hypothetical nodulating maize plant with the modeled RGR penalties, we calculated the Value Cost Ratio (VCR) of fertilizing maize with typical nitrogen fertilizer and the VCR of adding rhizobia inoculum with yield and price estimates for the United States (USA) and sub-Saharan Africa (SSA) **(Table 3)**.

**Table 3:** VCR and input parameter values comparing the profitability of fertilizer and rhizobia inoculum application versus using no fertilizer or inoculum amendments in the USA and SSA.

| Source | Parameter | Value | Condition | VCR |
| --- | --- | --- | --- | --- |
| USA | Fertilizer price (USD ha<sup>-1</sup>) | 198 (45) | + Fertilizer | 3.4 |
|  | Yield with fertilizer (tonne ha<sup>-1</sup>) | 11.5 (81) |  |  |
|  | Inoculum price (USD ha<sup>-1</sup>) | 9.9 (43) | + Inoculum | 47.0 |
|  | Yield with inoculum (tonne ha<sup>-1</sup>) | 10.0 (81)<sup>a</sup> |  |  |
|  | Corn price (USD tonne<sup>-1</sup>) | 141 (82) | Control |  |
|  | Control Yield (tonne ha<sup>-1</sup>) | 6.7 (83)<sup>b</sup> |  |  |
| SSA | Fertilizer price (USD ha<sup>-1</sup>) | 232 (46)<sup>c</sup> | + Fertilizer | 2.3 |
|  | Yield with fertilizer (tonne ha<sup>-1</sup>) | 3.6 (46) |  |  |
|  | Inoculum price (USD ha<sup>-1</sup>) | 38 (9,44) | + Inoculum | 11.2 |
|  | Yield with inoculum (tonne ha<sup>-1</sup>) | 3.1 (46)<sup>a</sup> |  |  |
|  | Corn price (USD tonne<sup>-1</sup>) | 230 (46) | Control |  |
|  | Control Yield (tonne ha<sup>-1</sup>) | 1.3 (46) |  |  |

<sup>a</sup>Yield with inoculum calculated by assuming that relative growth rate penalties translate 1:1 into yield penalties. We assume the worst-case scenario for the use of nitrogen-fixing maize by applying the largest relative growth rate penalty observed (12.8% penalty at seedling stage).
<sup>b</sup>Control yield value taken from a report on a plot that had not had fertilizer amended for 2 years (83).
<sup>c</sup>Taken from average fertilizer application needed to achieve optimal yield across a range of SSA sites in the mechanistic and empirical models presented in Bonilla-Cedrez 2021 (46)

Our results show that in both the USA and SSA, the potential profitability of a nitrogen-fixing maize crop is greater than that typically observed with regular nitrogen fertilizer additions, with a VCR of 45.4 vs 3.4 for the USA and a VCR of 11.2 vs 2.3 for SSA respectively. The greater potential profitability in the USA is driven by the much lower price of inoculum in the USA ($9.9 ha^-1^)(43) as compared with SSA ($38 ha^-1^)(9,44). This large difference in inoculum cost contrasts with the similar nitrogen fertilizer costs in the USA ($198 ha^-1^)(45) and SSA ($205 ha^-1^)(46).

### Addition of N_2_-fixing rhizobia symbiosis in maize would provide RGR benefits stabilized by the presence of AMF

To quantify the impact *Zea mays*’ existing AMF symbioses may have on the efficacy of engineering a novel symbiosis with an N_2_-fixing rhizobium, we constructed a model featuring maize in association with both symbionts and then compared the predicted RGR improvements from adding rhizobia to maize models with and without AMF (**Figure 3**).

**Figure 3:**
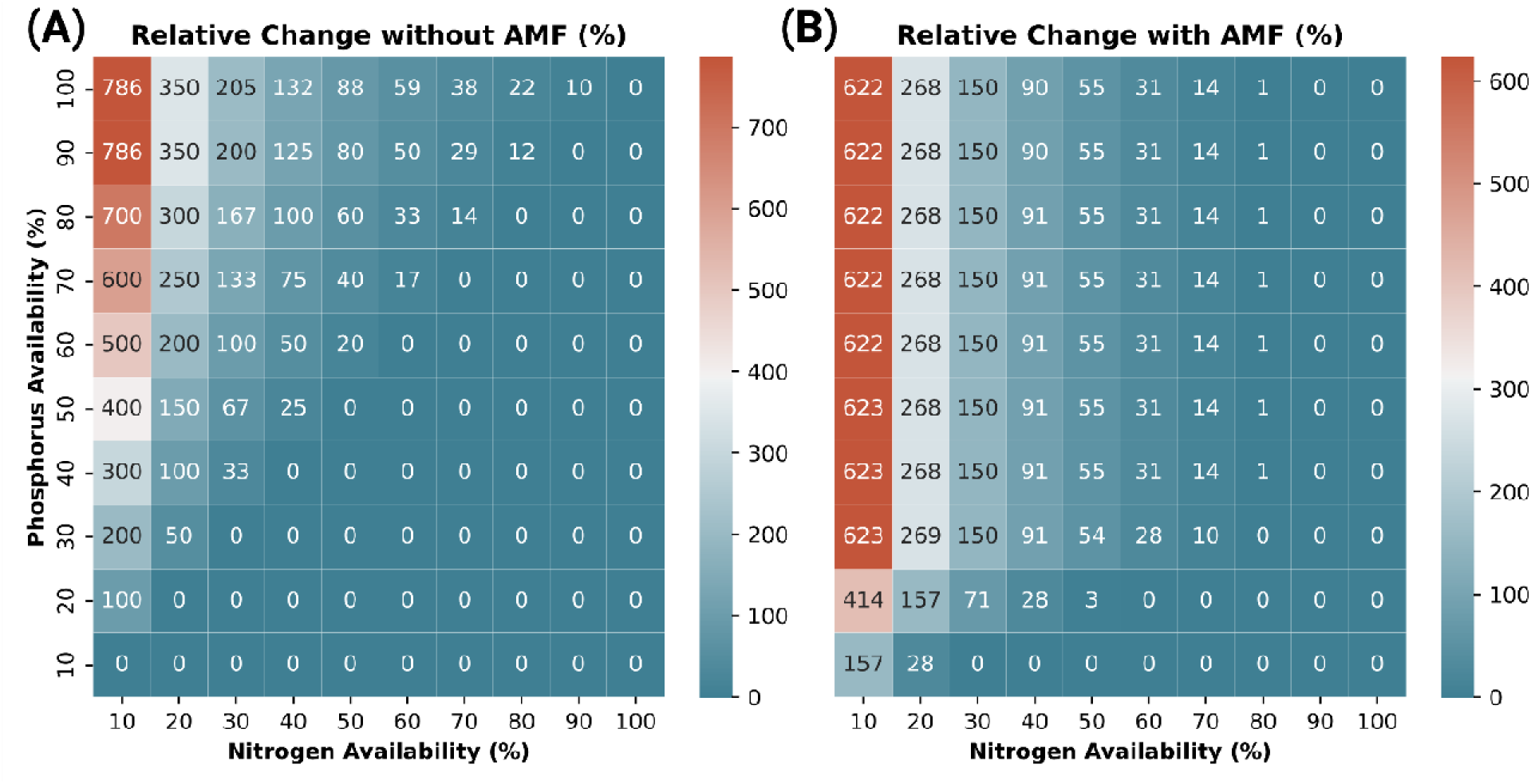
Comparison of predicted RGR improvement resulting from addition of N_2_-fixing rhizobia to *Z. mays* when AMF is absent. **(A)** and present **(B)**. Relative change in both cases is calculated by taking the difference in RGR between a base case and that same model with the rhizobia added – in **(A)** this is the difference between the *Z. mays* model alone and the *Z. mays* + *B. diazoefficiens* model, and in **(B)** this is the difference between the *Z. mays* + *R. irregularis* model and the *Z. mays* + *B. diazoefficiens* + *R. irregularis* model.

When AMF is not present, we the addition of rhizobia results in improved plant RGR, with benefits being most pronounced when external nitrogen is low **(Figure 3A-B)**. However, in order to realize these RGR benefits, sufficient external phosphorus is needed to support additional biomass accumulation from rhizobia-provided nitrogen. The dependency of rhizobia-mediated RGR benefits on phosphorus is greatly reduced when the AMF symbiosis is present **(Figure 3C-D)**, due to the dramatic P uptake benefits conferred by the AMF. This leads to more consistent gains in RGR across the profile of external N and P availability, with the differences most pronounced under extremely N-deprived and moderately P-deprived conditions. This observation of the presence/absence of one symbiont impacting the RGR benefits of the other led us to quantify the synergy – or antagonism – between rhizobia– and AMF-mediated growth benefits across different soil conditions.

### Rhizobia and AMF provide synergistic benefits under nutrient-poor conditions, but are antagonistic under nutrient-replete conditions

To better understand the conditions under which rhizobia and AMF-mediated nutrient benefits complement or conflict with one another, we predicted growth benefits across different phosphorus and nitrogen levels with the full maize-rhizobia-AMF model and compared them with the additive benefits of using each strategy in isolation. By calculating the difference between the RGR benefits predicted by the combined model and the additive predicted benefits from the individual plant-symbiote models (**Figure S3**), we were able to quantify the synergy and antagonism between these strategies as a function of nutrient availability.

Under phosphorus depletion and low to medium nitrogen availability our model predicts that the two symbiotic relationships work synergistically by relaxing the nutrient limitation imposed by both phosphorus and nitrogen and allowing the plant to achieve near-optimal RGR with less than 100% N and P availability. Maximal RGR improvement is observed under conditions of phosphorus depletion with low to moderate nitrogen availability (**Figure 4B**). In this new maximal region, the relative benefit increased from 23% using only AMF or 400% using only rhizobia to 789%, corresponding with an absolute increase of RGR to 0.076 day^-1^. The synergism between the symbiotic relationships is largest (409%) when nitrogen availability is at 10% and phosphorus levels are at 40% (**Figure 4C**). The predicted RGR of plants grown under these nutrient conditions with both the AMF and rhizobia symbiosis is 0.085 day^-1^, roughly double the value predicted assuming the rhizobia and AMF are non-synergistic (0.041 day^-1^).

**Figure 4:**
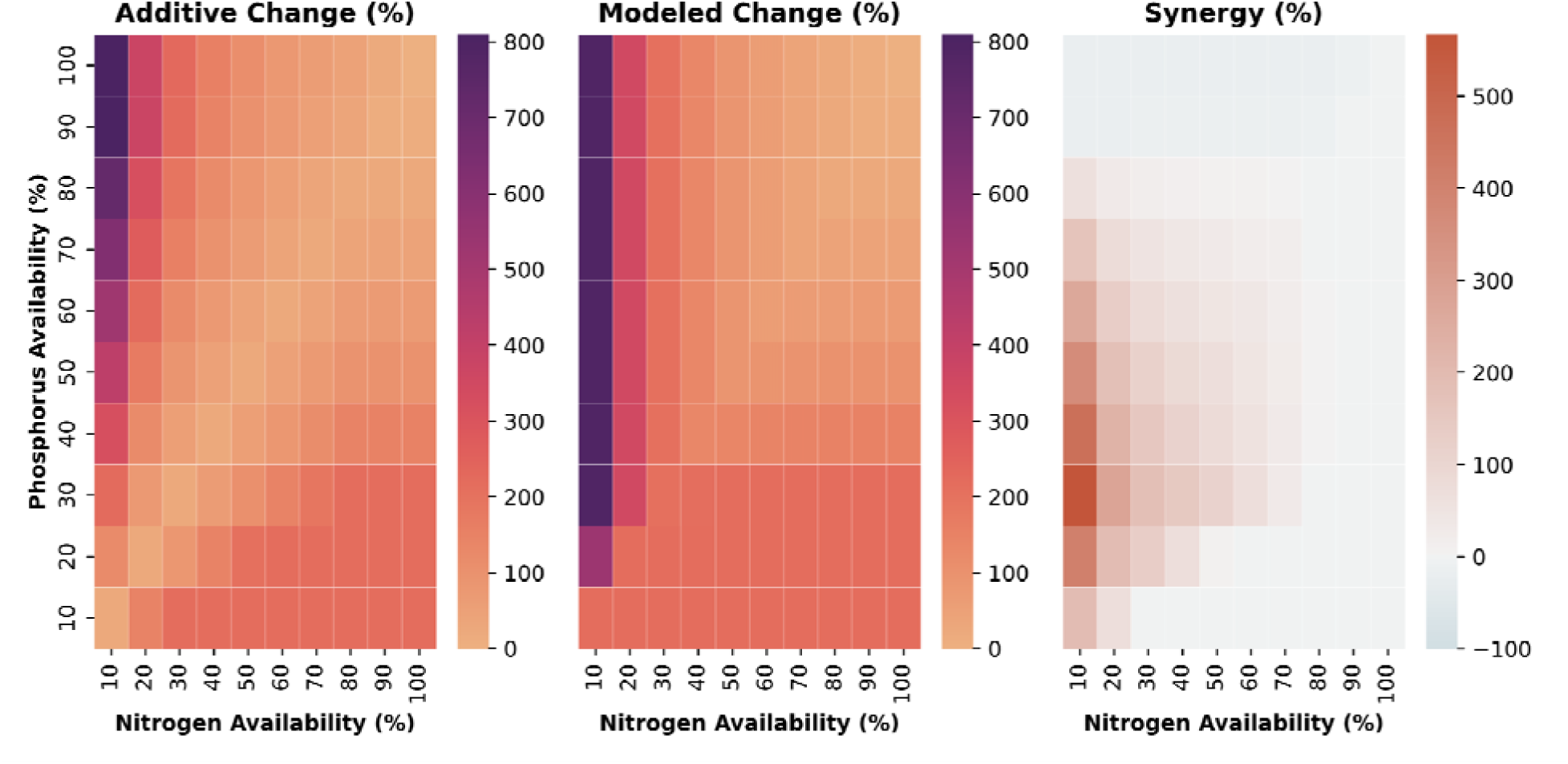
Additive and modeled RGR benefits and calculated synergies and antagonisms in the *Zea mays*, *Bradyrhizobium diazoefficiens*, *Rhizophagus irregularis* three-species system when the maize model’s biomass is representative of seedling C:N ratios. **(A)** Sum of relative growth rate changes from adding *B. diazoefficiens* or *R. irregularis* to the *Z.* mays model under given nitrogen and phosphorus availabilities. **(B)** Modeled relative growth rate changes in the three-species model. **(C)** Synergy and antagonism between *B. diazoefficiens* and *R. irregularis* quantified as the difference between the modeled and additive growth rate changes. For the additive change, modeled change, and synergy when the maize model’s biomass is representative of plants at the jointing or silking stages, refer to **Figures S7-8**.

Comparing this with the RGR predicted for plants without either symbiont but with 100% of their nutrient needs met (0.096 day^-1^) suggests that plants could achieve ∼89% of their maximum RGR with only 10% and 40% of optimal N and P availability. Under these conditions, the maize plants would get the majority of their N from their rhizobial symbionts (∼86%) and only (∼3%) from their AMF symbionts **(Dataset S2)**, consistent with the lower g C g N^-1^ associated with rhizobia versus AMF.

Maximum antagonism (–21%) is observed under phosphorus-replete conditions, whereas the model predicts negligible antagonism (less than –1%) under nitrogen-replete conditions. The magnitude of the positive synergy under moderate-to low-nitrogen availability and moderate phosphorus availability, expressed in terms of relative change, (48-409%) far outweighs the worst antagonisms seen (0-21%). However, the worst antagonisms are observed under phosphorus-replete conditions, some of which support high basal RGR. This causes the worst antagonism in terms of absolute RGR differences under phosphorus-replete conditions (–0.016 day^-1^) to be closer to the highest positive synergy values (0.045 day^-1^) than one would expect based purely on the relative changes.

## Discussion

### Modeling the benefits of AMF on plant nutrition and growth

To our knowledge, this study presents the first metabolic model describing the exchange of carbon, nitrogen, and phosphorus between a plant and AMF. Our plant-AMF framework reproduced empirically observed RGR benefits and predicted carbon investments consistent with prior literature (40), and represents a promising platform for computational investigations into plant-AMF symbiosis. This framework could be further strengthened with additional information on below-ground AMF biomass quantities and the interdependence of nitrogen uptake, phosphorus uptake, and plant biomass gains. The data currently available is limited and has widely varying estimates of the impacts of AMF, particularly regarding AMF-mediated nitrogen uptake (11,47,48).

### Impact of existing AMF symbiosis on potential benefits of rhizobia engineered into maize

Given the potential benefits of adding rhizobia to cereal crops and the possible interactions of existing AMF-associations with this engineered symbiosis, the question of whether these symbioses complement or hinder each other is crucial. Previous work has shown that rhizobia and AMF can enhance each other’s colonization of plant tissues and improve plant growth synergistically (15). Our results corroborate these findings and provide quantitative predictions of the magnitude of these effects and nutrient conditions under which they are optimized. In contrast to a plant without an AMF symbiosis, a maize plant with this symbiosis is predicted to achieve substantial and consistent growth benefits across a wider range of soil nutrient conditions. We predict that synergy between the AMF and rhizobia strategies roughly doubles the relative growth rate of maize plants under severe N deprivation and moderate P availability compared to a scenario where no synergy between these strategies exists. We also predict significant antagonism between these strategies under phosphorus-replete conditions. These findings suggest that existing AMF symbioses are likely to enhance the benefits of a hypothetical rhizobia symbiosis in cereal crops.

### Viability of engineering N_2_-fixing rhizobia symbiosis into maize

Both the addition of rhizobia to cereal crops and the enhancement of AMF-associations and accompanying nutrient-uptake benefits are strategies being pursued to sustainably improve agricultural production (4,12,49), so the models presented in this study describing the maize-rhizobia and maize-AMF symbioses in isolation are also important to consider. We explored the efficiency of engineering rhizobia N_2_-fixation and enhanced AMF association individually. We predict a lower RGR penalty (3.6 – 12.8%) than previous work (∼14%) that predicted the penalty in maize by comparing the overall (grain + stover) N content of soybean (3.1%) and maize (1.4%) at harvest (9,50,51). We also find that the RGR penalty is highest earlier in vegetative growth when the nitrogen content of plant tissue is higher, so focusing on end-point measurements of plant biomass composition may not give the best estimate of the cost of nitrogen fixation over a growing season. However, our results support the idea that cereal crops should experience smaller RGR losses than legumes when relying on N_2_-fixation **(Table 2),** making this an attractive bioengineering approach to sustainably improve cereal crop yields.

Our VCR analysis (**Table 3**) indicates that inoculation of a nitrogen-fixing maize plant could be a profitable alternative to the use of nitrogen fertilizer in both the USA and SSA. However, the profitability would be greater in the USA due to the almost fourfold greater price of inoculum on a per hectare basis in SSA as compared with the USA (9,46). This suggests that, if nitrogen-fixing maize is developed, reducing the cost of inoculum for cultivators will be important for driving the adoption of this technology in SSA. One caveat to this analysis is the great deal of variability in maize yield potential seen across SSA, with some regions having much higher potential yields upon fertilization, closer to 7 tonne ha^-1^ as compared with the 3.6 tonne ha^-1^ used in our analysis (46). Another caveat is that the degree to which inoculation with rhizobia is necessary to achieve good levels of biological nitrogen fixation varies with the natural abundance of the particular species. Here, we have modeled *Zea mays* in a hypothetical association with *Bradyrhizobium diazoefficiens*, but other species could require less inoculum per hectare, making the cost of inoculum less influential in the cost-benefit analysis.

### Limitations and Future Directions

A major limitation of the models presented in this study is the fact that only metabolic interactions are considered. While a model incorporating all of the diverse regulatory details and potential interactions between plants, rhizobia, and AMF would not be possible given our current knowledge of these systems, it should be noted that unmodeled factors, such as water-use impacts or regulatory cross-talk between rhizobia and AMF, may greatly impact the real world system. Additionally, there is room to expand on the models presented in this study to further investigate the costs and benefits of symbiosis on maize. Although we have made an effort to partially explore the impact of the plant’s developmental stage on the costs and benefits of the rhizobia symbiosis, a quantitative model describing how symbiont presence/absence and soil nutrient levels affect biomass composition would greatly enhance the predictive capabilities of the presented metabolic models. Additionally, the *R. irregularis* model could be expanded to account for the different tissue types seen in that organism (23). Further, in our framework, the fungus is treated as carbon-limited and does not compete for external nutrients with the plant, so the parasitism predicted by the trade-balance model (13) under nutrient-replete conditions is not predicted by our model. Explicitly modeling this resource competition could address this.

Finally, to quantify the impact of these symbioses on grain production, a developing kernel tissue compartment could be added to the maize model, and incorporating the dynamic changes to RGR corresponding to the different developmental stages with a crop growth model would allow us to translate RGR changes into yield predictions. Indeed, in light of interest in developing maize lines with modified kernel N-content and a growing season optimized to better make use of soil N, a modeling framework that integrates detailed flux maps with yield potential in the context of symbioses that affect plant nutrition could be very valuable (52).

### Conclusion

This work presents the first analysis of the predicted metabolic flux costs and benefits of engineering N_2_-fixation in a cereal crop by rhizobia alone and together with AMF. Our results support previous predictions that cereals represent an attractive target for engineering to associate with rhizobia due to their lower nitrogen content. We also developed the first experimentally validated metabolic model describing the exchange of carbon, nitrogen, and phosphorus between plants and AMF. Creating a combined plant-AMF-rhizobia model, we predicted substantial synergy between the two symbioses when nitrogen and phosphorus availability is low and antagonism when phosphorus availability is adequate. Our results indicate that these engineering strategies could efficiently and economically improve cereal crop yields

## Materials and Methods

### Maize shoot/root model construction

A bottom-up reconstruction of *Arabidopsis thaliana’s* metabolism (22) was used as a base representation of plant central metabolic processes. As done in a previous C4 metabolic modeling study (53), this model was duplicated to create separate bundle sheath (BS) and mesophyll (M) models. Other additions and refinements of the bottom-up reconstruction (22) described in the C4 modeling study (53) were also made to improve the accuracy of the model, with the exception of the latter’s modifications to the chloroplast ATP synthase reaction, since the original model already correctly captured the c-subunit stoichiometry and consequent ATP/proton stoichiometry typical of plant systems (54).

These models were then connected by transport reactions representing exchanges between BS and M cells through the plasmodesmata and flux through cytosolic phosphoenolpyruvate carboxylase and mitochondrial malate dehydrogenase 1 were disallowed, based on the methods described in (53) to ensure that the flux map of the resulting model lines up with canonical carbon assimilatory fluxes through the NADP-ME subtype characteristic of maize (55). As in the earlier soybean-rhizobium FBA study (9), and based on earlier literature derivations (56), an upper bound of 2920 µmol g^-1^ DW h^-1^ on photon uptake was imposed.

Another duplicate of the central metabolic model (22) was then made to represent the plant’s root tissue. Metabolite exchanges were adopted from (20). All metabolite exchanges between the shoot tissue model and the environment, apart from CO_2_, O_2_, and light, were disallowed. The root tissue model was treated the same, except with the additional capacity to take up ammonium, orthophosphate, water, and dihydrogen sulfide. The orthophosphate uptake reaction was modified based on the fact that direct phosphate uptake is thought to occur through proton symporters (57). Two protons were assumed to move across the cell membrane for each phosphate (58).

Base biomass compositions for the root and shoot tissue were adapted from (27), with additional reactions from KEGG (59) added to produce biomass components not covered by the original *Arabidopsis* central metabolism model (22). A list of added reactions can be found in **Dataset S1**. The ratio of shoot to root biomass was constrained to a value of 90:10 (60).

Growth-associated maintenance costs (GAM) and non-growth associated maintenance costs (NGAM) for maize were calculated from literature data (61) and modeled either as generic ATP or glucose burning at a specified rate **(Dataset S4)**. Growth-associated maintenance was calculated as 19 mmol ATP gDW^-1^ d^-1^, lower than the assumed 30 ATP gDW^-1^ d^-1^ featured in many other plant metabolic models (62). The NGAM cost was calculated as 5.76 mmol ATP gDW^-1^ d^-1^. All maintenance costs were distributed across tissues according to their proportions of total dry weight.

### AMF maize model construction and analysis

We used the previously published and validated GEM of the AMF *Rhizophagus irregularis* to represent the fungal partner in a plant-fungus symbiotic relationship (23). To make the *R. irregularis* model dependent on the plant for its carbon and energy needs, all external uptakes were disabled except for those with oxygen, carbon dioxide, dihydrogen sulfide, ammonium, phosphate, and a number of mineral micronutrients necessary for biomass accumulation. Based on literature evidence, only a small number of metabolites were allowed to be exchanged between the fungal and plant root models (57) **(Dataset S5**). From the plant, sugar (in the form of glucose) and palmitic acid were allowed to be transported to the AMF. From the fungus, phosphate and ammonium were allowed to be transported, with a cost of 0.5 ATP per orthophosphate transported (58).

### Recalculation of biomass coefficients to account for changing tissue C:N ratios

Published C:N ratio data (63) was used by modifying the base maize biomass composition from Saha et al. 2011 (27) to match the measured shoot and root C:N ratios. The nitrogenous biomass (NB) and non-nitrogenous components of the biomass (B) were grouped together and the appropriate proportion for these groups were calculated as follows:

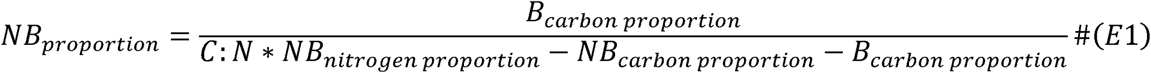

Where *B_carbon proportion_* is the proportion of the weight of the non-nitrogenous biomass that is made up of carbon, *NB_nitrogen proportion_* and *NB_carbon proportion_* are the proportions of the weight of the nitrogenous biomass that is made up of nitrogen and carbon respectively, and *C:N* is the desired C:N ratio. We then split the proportion of non-nitrogenous biomass into the individual non-nitrogenous biomass components in a way that maintains their ratios relative to each other from the original biomass equation (27). Finally, these weight proportions were divided by the average molecular weight of each class of compounds to give the coefficients for each in the final biomass equation in units of mmol gDW^-1^ day^-1^.

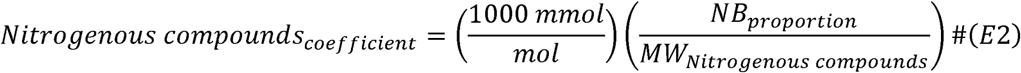

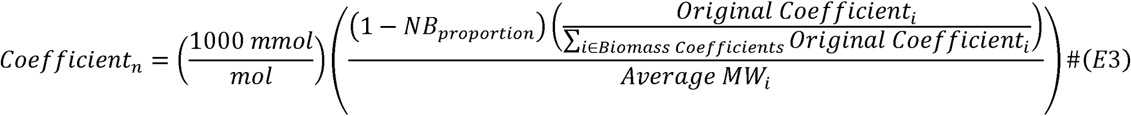

Where *Original Coefficient_i_* is the coefficient for a given biomass component *i* in the original biomass equation (27), *Biomass Coefficients* is the set of all of these original coefficients, and *Average MW_i_* is the average molecular weight of the constituent metabolites in a group of biomass compounds. Note that the final coefficients from this calculation result in biomass equations whose components add up to 1 gDW.

### Nodulated maize model construction and analysis

The workflow previously used in (9) was adapted to create a multi-species model with the maize C4 root/shoot model described above as the plant component and the GEM of *Bradyrhizobium diazoefficiens* as the N_2_-fixing bacterial symbiont (24).

First, a duplicate of the root model, representing the tissue of a hypothetical maize nodule, was made with exchanges between it and the root limited as in Holland et al. 2023 (9) and as listed in **Dataset S5**. Next, transporters describing the exchange of dicarboxylates and ammonium were added between the nodule model and the *B. diazoefficiens* model to reflect known exchanges from the literature (9). Allantoate is thought to be the primary form of nitrogen exchanged between nodule tissue and the plant root, so reactions describing allantoate biosynthesis and degradation were added to the model from KEGG (59).

The non-growth associated maintenance (NGAM) costs of *B. diazoefficiens* were assumed to be the same as those for the *Medicago truncatula* bacteroid *Sinorhizobium meliloti*: 2.52 mmol gDW^-1^ h^-1^ (64). Assuming that the bacteroid constitutes 2% of the biomass of the overall system (9), this value was scaled to 1.21 mmol gDW^-1^ d^-1^. At steady-state we assume that the bacteroid is not accumulating biomass, so growth-associated maintenance costs were ignored. Note that the *M. truncatula / S. meliloti* symbiosis features indeterminate nodule formation and amide export, in contrast to the determinate nodules and ureide export of the *Glycine max / B. diazoefficiens* symbiosis we are elsewhere modeling the hypothetical *Z. mays* / *B. diazoefficiens* symbiosis after. These differences likely result in differences in NGAM; however, as shown in **Figure S2**, model outputs are relatively robust to changes in NGAM and associated parameters, like tissue proportions, suggesting that small variations in these costs will not have a large impact on study conclusions.

### Optimization details

Flux Balance Analysis optimizations were performed, in most simulations, to maximize plant RGR.

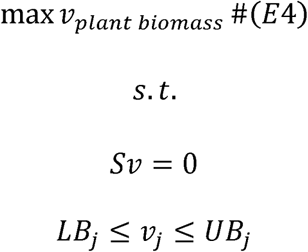

Where *S*, represents the stoichiometric matrix of the metabolic network model being optimized, *v* is the vector of all fluxes in that network, and *LB* and *UB* are vectors of the lower and upper bound constraints. In this study, all optimizations were followed up by minimization of the sum of fluxes in the network via regularization of the L1-norm (65), often referred to as parsimonious FBA (pFBA) (66), which corresponds to the following optimization:

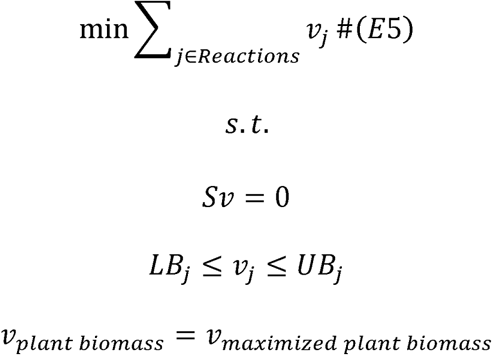

Where *j* is the index of a reaction in the list of all reactions in the network (*Reactions*) and *v*_maximized plant biomass_ is the maximized biomass value obtained from the initial optimization.

### Model optimization and analysis

Growth benefits associated with N_2_-fixation, AMF association, and both N_2_-fixation and AMF association in concert, were estimated by comparison of predicted plant RGR in the base maize model and the appropriate symbiosis models across a range of soil nitrogen and phosphorus uptake rates **(Figure S2)**. All nitrogen uptake by the plant and, in models that feature it, the AMF symbiont is assumed to be in the form of ammonium.

Optimizations of the maize model with an added N_2_-fixing bacterial symbiont were performed as in Holland et al. 2023 (9). Briefly, the plant model without the bacterial symbiont is first optimized to maximize RGR and the ammonium uptake necessary to support this maximized growth rate is recorded. This ammonium uptake rate defines the point at which the model’s biomass accumulation is no longer limited by ammonium availability. For both the model with and without the associated N_2-_fixing symbiont, we then continuously decrease the external ammonium level and evaluate the RGR at each point. We also calculate the “grams of carbon per gram nitrogen,” or g C g N^-1^, which is a measure of the ratio of carbon, in the form of glucose or fatty acids, sent to the fungus from the plant in exchange for nitrogen from the fungus. g C g N^-1^ was calculated as in Holland et al. 2023 (9):

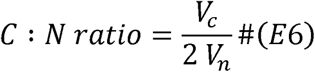

Where V_c_ is the rate of CO_2_ release from the nodule tissue and V_n_ is the flux through the bacteroid nitrogenase reaction.

### AMF model optimization and analysis

The AMF model was optimized in a sequential process meant to mathematically encode the following assumptions about the plant/AMF symbiosis:

1. Since the AMF fungus is dependent on carbon from the host plant, the ability of the fungus to confer nutrient uptake benefits to the plant is a function of carbon investment from the plant.
2. To achieve a measured level of nutrient uptake benefits because of AMF inoculation, the modeled system must achieve the same ratio of AMF to plant biomass, with decreasing ratios linearly decreasing the nutrient uptake benefits conferred.

The data for phosphorus uptake benefits in AMF– and AMF+ plants (37), data for nitrogen uptake benefits (38), and hyphal length density (HLD)(37) values were used to estimate AMF biomass necessary to achieve maximal nutrient uptake benefits. We estimated the dry weight of AMF tissue (*DW_AMF_*) from the HLD values and assuming the plants were grown in 2,750 grams of soil (37), as described in Equation E7.

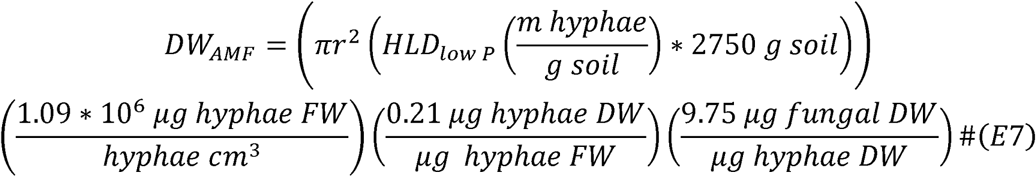

Where *HLD*_low P_ is the hyphal length density in centimeters reported in the lowest phosphorus condition. The conversion between hyphae fresh weight (FW) and hyphae volume, 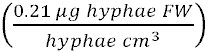, is taken from literature estimates, as is the fresh weight to dry weight (DW) conversion (39). Finally, given that substantial fungal biomass is contained in non-hyphal tissue like vesicles, we used a reported ratio of hyphal to total (hyphal plus vesicle) biomass (35), 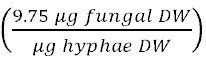 to convert from the hyphal biomass estimate to an estimate of total fungal dry mass.

To optimize the system, the maximal biomass growth of the system without any AMF-mediated nutrient uptake benefits is calculated across a range of phosphate and ammonium uptake rates as described in **Eq. 4 – 5**.

The ratio of the AMF-plants’ biomass and the AMF itself in the low-phosphorus condition reported in Sawers *et al.* 2017 (37) was then used to define the following reactions in the model:

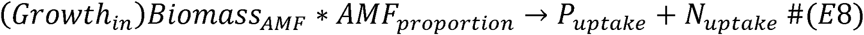

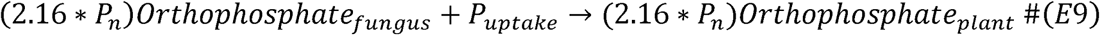

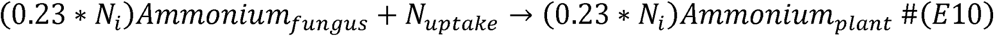

Where *Growth_in_* is the maximal biomass accumulation that the plant is capable of in the absence of AMF-mediated nutrient uptake benefits, AMF_proportion_ is the proportion of AMF-to-plant biomass, and *P_n_*and *N_i_* are the non-AMF-mediated plant phosphate and ammonium uptake rates. A single unit of flux through this reaction requires producing the observed amount of fungal biomass relative to plant biomass, yielding a single unit of *P_uptake_* and *N_uptake_*, which can be used in **Eq. 9** and **Eq. 10** to move 2.16 times the basal phosphate uptake rate or 0.23 times the basal ammonium uptake rate. The upper bound of **Eq. 8** is set to 1, such that no more than 2.16 times the basal phosphate uptake rate and 0.23 times the ammonium uptake rate can be achieved, based off of reported nitrogen and phosphate uptake benefits (37,38).

With these additions, the full model, plant and AMF partner, is optimized as described in **Eq. 4-5.** In addition to the maximal RGR of the plant, the fluxes of glucose and palmitate from the plant to the fungus and the net CO_2_ uptake of the plant are recorded. From these quantities, we calculate the percent of CO_2_ from the plant that is invested in the AMF partner:

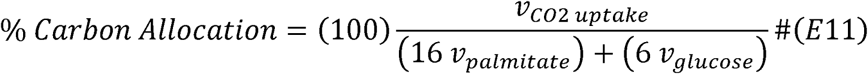

Where *v_CO2 uptake_* is the plant’s CO2 uptake flux and *v_palmitate_*and *v_glucose_* are the fluxes of palmitate and glucose transport from the plant to the fungus.

### AMF growth rate validation

To generate AMF-incompatible (AMF-) maize hybrids for field evaluation, a B73 inbred line homozygous for the *castor-2* mutation (67) was crossed as male to a W22 inbred line also homozygous for *castor-2*. Control AMF-compatible (AMF+) hybrids were generated by crossing the same B73 *castor-2* line as male to an AMF-W22 inbred line carrying a mutation in the maize ortholog of the *pollux* gene (41,68), with trans-complementation recovering AMF compatibility in the F_1_ hybrid progeny. This scheme allowed that both AMF– and AMF+ hybrids were produced from AMF-mothers to reduce any strong maternal effect.

AMF+ and AMF-maize hybrids were evaluated at Russell E. Larson Agricultural Research Center at the Pennsylvania State University (40°42’39.1”N 77°57’11.4”W) within a larger split-split-plot design with 2.5-foot row spacing receiving drip irrigation. Prior to planting, the soil of 90 random rows (14.1 m^2^ density) was sampled by pooling soil from the front, middle, and end of rows at a depth of 0-10 cm. Soil samples were then submitted to the Agricultural Analytical Services Lab to determine Total N and Mehlich 3 extractable P by ICP analysis with soil parameters at unsampled rows modeled with kriging interpolation using the ‘gstat’ R package(69). Seeds were planted on May 18^th^, 2023, one week after 150 lbs of 19-19-19 (%N-%P-%K) fertilizer was applied. After receiving an additional 300 lbs of urea per acre on June 21^st^, 2023, AMF+ and AMF-plant biomass was sampled, dried, ground, and analyzed with a dry combustion elemental analyzer to determine Total N content (as described in (70)) at timepoints T1-T4 (28, 38, 48, and 58 days after emergence) and at harvest on October 4, 2023. To assay AMF colonization in AMF+ (32% colonization; n=117) and AMF-(2% average colonization, n = 57) plants, root tissue was harvested at 8-weeks after emergence to quantify fungal colonization via microscopy following staining. Briefly, ∼0.25 g of fine root tissue was collected from 0-0.25 m depth, cleared in 10% KOH at 70°C over 48 hours, rinsed five times in deionized water, incubated in 0.3 M HCl for 15 mins at room temperature, and stained at 96°C for 8 min using a 0.05% w/v Direct Blue 15 Dye staining solution in a 1:1:1 mixture of lactic acid: glycerol: deionized water. Percent root length colonized was then determined by noting the presence and absence of AMF structures at 40x magnification along 100 intersections on a 1 cm x 1 cm grid (71).

Available N in the field trial was estimated by adding the known N application rate to an estimate of N mineralization from soil organic matter. Soil organic matter (SOM) content was estimated from measured total N, assuming a standard composition of SOM of 5% nitrogen (72,73). From this, we calculate the contribution of mineralized organic nitrogen as:

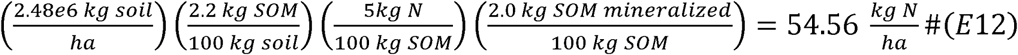

Where the 2.2 kg SOM 100 kg^-1^ soil is derived from the measured total N, the weight of a hectare of soil is estimated assuming a bulk density of 1.24 g/cm^3^ in the 0-20cm depth (74), SOM is assumed to be 5% N by weight (72,73) and 2.0 kg of SOM is assumed to be mineralized per 100 kg of SOM per growing season (75). The 0-20cm depth is used for the mineralization calculation because the majority (∼80%) of the root surface area of a maize plant is present between soil depths of 0-20cm (76).

The RGR of plants from the field trial dataset was estimated using the following equation:

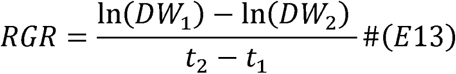

Where *DW_1_*and *DW_2_* are the shoot dry weight measurements (g) at growth timepoints 1 (*t_1_*) and timepoint 2 (*t_2_*), which are 28 and 38 days after emergence, respectively. These two early growth timepoints were used to calculate RGR to make for as direct a comparison between the AMF– and AMF+ samples as possible by minimizing any potential impact of differing developmental trajectories between AMF+ and AMF – plants.

The average soil phosphorus level across all planted rows was approximately 59 mg kg^-1^ and the average total nitrogen content was 0.11% (kg N kg^-1^ soil) **(Dataset S3).** A previous study (37) showed that both shoot dry weight gains and AMF-mediated growth benefits were previously shown to disappear somewhere between phosphorus concentrations of 15.5 mg kg^-1^ and 53.2 mg kg^-1^. This indicates that the plants in this study had adequate phosphorus available to achieve optimal or near-optimal growth. Corroborating this idea, one study found that high-performing maize lines take up an average of 114 kg P_2_O_5_, or 50 kg P, during the growing season (77), as compared to the 66 kg P that were available to a depth of 20 cm in the samples with the lowest soil P concentration in this study (26.5 mg/kg). This means that the plants assessed in this study received 100%+ of the P necessary to achieve optimal growth. The average total nitrogen content in our plots of 0.11% was measured prior to an application of 187 kg N ha^-1^. Together, the mineralization of the soil organic matter and the applied N fertilizer amount to the equivalent of an input of 241 kg N ha^-1^. Assuming that our maize hybrids can achieve similar optimal growth rates as those reported in a previous study assessing optimal N application rates and nutrient uptake rates (77), high-performing maize hybrids will take up an average of 286 kg N ha^-1^. This means that the plants assessed in this study received ∼84% of the N necessary to achieve optimal growth. Together with the P availability, this makes the nitrogen-limiting upper-right-hand quadrant the appropriate comparison between the RGR improvements in **Figure 1A** and our empirical measurements.

### Sensitivity analysis

Local sensitivity analysis was performed by varying the values of key input parameters for all models presented in this study by ±10% and then calculating the percent change in (i) RGR, (ii) g C g N^-1^, and (iii) g C g P^-1^. The reported sensitivity values are the averages of the percent changes resulting from increasing and decreasing a given parameter value.

### Monte Carlo estimation of uncertainty

A Monte Carlo approach was implemented to calculate the distributions of RGR benefits and carbon allocation percentages from the plant-AMF model. The uncertainty of the AMF-mediated P uptake benefits and the necessary AMF biomass was estimated from (37,39). Uncertainty data was not available for the AMF-mediated N uptake benefits, so the coefficient of variation of this parameter was assumed to be the same as that of the P uptake benefit. 1,000 Monte Carlo samples were then generated by varying these three parameters according to their means and standard deviations **(Table S1)** and the resulting RGR and carbon allocation estimates were used to generate confidence intervals.

### Software

Optimizations and model analysis were performed in MATLAB R2020a using the COBRA Toolbox 3.0 (78), the Enterprise IBM ILOG CPLEX solver version 12.10, and the Gurobi Optimizer 11.02. Code to run model and reproduce manuscript results can be found at: https://github.com/Matthews-Research-Group/Maize-Rhizobia-AMF_FBA.

## Acknowledgments

This work was supported by the project Enabling Nutrient Symbioses in Agriculture (ENSA), that is funded by Bill & Melinda Gates Agricultural Innovations (INV-57461) (JAMK, RJ, PS, RJHS, and MLM). PS and RJHS were supported by the United States Department of Agriculture (USDA) project *The genetic basis of maize response to arbuscular mycorrhizal fungi* (2022-67013-38264). RJHS is supported by the USDA National Institute of Food and Agriculture and Hatch Appropriations under Project #PEN04734 and Accession #1021929. Figures made using <u>BioRender.com</u>.

## Data availability

All data, models, and code needed to reproduce the analysis in this study can be found in the online supplemental material accompanying the manuscript and at https://github.com/Matthews-Research-Group/Maize-Rhizobia-AMF_FBA.

## Author contributions

JAMK and MLM conceived the study. JAMK built, made predictions with, and analyzed the results of the metabolic models presented in this study. RJ performed the sensitivity analysis. PS and RJHS generated and evaluated AMF compatible and incompatible maize hybrids. JAMK, RJ, and MLM interpreted the results. JAMK wrote the manuscript. All authors revised the manuscript and approved the final version.

## References

1. Gates B. How to avoid a climate disaster: the solutions we have and the breakthroughs we need. Vintage; 2021.

2. Boerner LK. Industrial ammonia production emits more CO2 than any other chemical-making reaction. Chemists want to change that. Chem Eng News. 2019;97(24):1–9.

3. Liu X, Elgowainy A, Wang M. Life cycle energy use and greenhouse gas emissions of ammonia production from renewable resources and industrial by-products. Green Chemistry. 2020;22(17):5751–61.

4. Jhu MY, Oldroyd GED. Dancing to a different tune, can we switch from chemical to biological nitrogen fixation for sustainable food security? PLOS Biology. 2023 Mar 14;21(3):e3001982.

5. Goodman PS, O’Reilly F. How a Fertilizer Shortage Is Spreading Desperate Hunger. The New York Times https://www.nytimes.com/2023/10/15/business/nigeria-fertilizer-shortage.html. 2023;

6. Kassouri Y. Fertilizer prices and deforestation in Africa. Food Policy. 2024 July 1;126:102674.

7. Santi C, Bogusz D, Franche C. Biological nitrogen fixation in non-legume plants. Annals of Botany. 2013 May 1;111(5):743–67.

8. Guo K, Yang J, Yu N, Luo L, Wang E. Biological nitrogen fixation in cereal crops: Progress, strategies, and perspectives. Plant Communications. 2023 Mar 13;4(2):100499.

9. Holland BL, Matthews ML, Bota P, Sweetlove LJ, Long SP, diCenzo GC. A genome-scale metabolic reconstruction of soybean and Bradyrhizobium diazoefficiens reveals the cost– benefit of nitrogen fixation. New Phytologist. 2023;240(2):744–56.

10. Smith SE, Read DJ. Mycorrhizal symbiosis. Academic press; 2010.

11. Beslemes D, Tigka E, Roussis I, Kakabouki I, Mavroeidis A, Vlachostergios D. Effect of Arbuscular Mycorrhizal Fungi on Nitrogen and Phosphorus Uptake Efficiency and Crop Productivity of Two-Rowed Barley under Different Crop Production Systems. Plants [Internet]. 2023;12(9). Available from: https://www.mdpi.com/2223-7747/12/9/1908

12. Hornstein ED, Sederoff H. Back to the future: re-engineering the evolutionarily lost arbuscular mycorrhiza host trait to improve climate resilience for agriculture. Critical Reviews in Plant Sciences. 2024;43(1):1–33.

13. Johnson NC. Resource stoichiometry elucidates the structure and function of arbuscular mycorrhizas across scales. New Phytologist. 2010 Feb 1;185(3):631–47.

14. Hoeksema JD, Chaudhary VB, Gehring CA, Johnson NC, Karst J, Koide RT, et al. A meta-analysis of context-dependency in plant response to inoculation with mycorrhizal fungi. Ecology Letters. 2010 Mar 1;13(3):394–407.

15. Liu XQ, Xie MM, Hashem A, Abd-Allah EF, Wu QS. Arbuscular mycorrhizal fungi and rhizobia synergistically promote root colonization, plant growth, and nitrogen acquisition. Plant Growth Regulation. 2023 July 1;100(3):691–701.

16. Afkhami ME, Stinchcombe JR. Multiple mutualist effects on genomewide expression in the tripartite association between Medicago truncatula, nitrogen-fixing bacteria and mycorrhizal fungi. Molecular Ecology. 2016 Oct 1;25(19):4946–62.

17. Schnepf A, Roose T. Modelling the contribution of arbuscular mycorrhizal fungi to plant phosphate uptake. New Phytologist. 2006 Aug 1;171(3):669–82.

18. Schnepf A, Leitner D, Schweiger PF, Scholl P, Jansa J. L-System model for the growth of arbuscular mycorrhizal fungi, both within and outside of their host roots. Journal of The Royal Society Interface. 2016 Apr 1;13(117):20160129.

19. de Vries J, Evers JB, Kuyper TW, van Ruijven J, Mommer L. Mycorrhizal associations change root functionality: a 3D modelling study on competitive interactions between plants for light and nutrients. New Phytologist. 2021 Aug 1;231(3):1171–82.

20. Chowdhury NB, Simons-Senftle M, Decouard B, Quillere I, Rigault M, Sajeevan KA, et al. A multi-organ maize metabolic model connects temperature stress with energy production and reducing power generation. iScience. 2023 Dec 15;26(12):108400.

21. Decouard B, Chowdhury NB, Saou A, Rigault M, Quilleré I, Sapir T, et al. Maize (Zea maysL.) interaction with the arbuscular mycorrhizal fungus Rhizophagus irregularisallows mitigation of nitrogen deficiency stress: physiological and molecular characterization. 2024;

22. Arnold A, Nikoloski Z. Bottom-up Metabolic Reconstruction of Arabidopsis and Its Application to Determining the Metabolic Costs of Enzyme Production. Plant Physiol. 2014 July;165(3):1380–91.

23. Wendering Philipp, Nikoloski Zoran. Genome-Scale Modeling Specifies the Metabolic Capabilities of Rhizophagus irregularis. mSystems. 2022 Jan 25;7(1):e01216–21.

24. Yang Y, Hu XP, Ma BG. Construction and simulation of the Bradyrhizobium diazoefficiens USDA110 metabolic network: a comparison between free-living and symbiotic states. Mol BioSyst. 2017;13(3):607–20.

25. Chowdhury NB, Simons-Senftle M, Decouard B, Quillere I, Rigault M, Sajeevan KA, et al. A multi-organ maize metabolic model connects temperature stress with energy production and reducing power generation. iScience. 2023 Dec 15;26(12):108400.

26. Dal’Molin CG de O, Quek LE, Palfreyman RW, Brumbley SM, Nielsen LK. C4GEM, a genome-scale metabolic model to study C4 plant metabolism. Plant Physiol. 2010 Dec;154(4):1871–85.

27. Saha R, Suthers PF, Maranas CD. Zea mays iRS1563: a comprehensive genome-scale metabolic reconstruction of maize metabolism. PLoS One. 2011;6(7):e21784.

28. Seaver SMD, Bradbury LMT, Frelin O, Zarecki R, Ruppin E, Hanson AD, et al. Improved evidence-based genome-scale metabolic models for maize leaf, embryo, and endosperm. Front Plant Sci. 2015;6:142.

29. Ehleringer J, Björkman O. Quantum Yields for CO2 Uptake in C3 and C4 Plants: Dependence on Temperature, CO2, and O2 Concentration 1. Plant Physiology. 1977 Jan 1;59(1):86–90.

30. Langdale JA. C4 cycles: past, present, and future research on C4 photosynthesis. Plant Cell. 2011 Nov;23(11):3879–92.

31. Ermakova M, Woodford R, Fitzpatrick D, Nix SJ, Zwahlen SM, Farquhar GD, et al. Chloroplast NADH dehydrogenase-like complex-mediated cyclic electron flow is the main electron transport route in C4 bundle sheath cells. New Phytologist [Internet]. 2024 July 22 [cited 2024 Aug 20];n/a(n/a). Available from: 10.1111/nph.19982

32. Pace BA, Perales HR, Gonzalez-Maldonado N, Mercer KL. Physiological traits contribute to growth and adaptation of Mexican maize landraces. PLOS ONE. 2024 Feb 1;19(2):e0290815.

33. Bago B, Pfeffer PE, Shachar-Hill Y. Carbon Metabolism and Transport in Arbuscular Mycorrhizas. Plant Physiology. 2000 Nov 1;124(3):949–58.

34. Wipf D, Krajinski F, van Tuinen D, Recorbet G, Courty PE. Trading on the arbuscular mycorrhiza market: from arbuscules to common mycorrhizal networks. New Phytologist. 2019 Aug 1;223(3):1127–42.

35. Olsson PA, Johansen A. Lipid and fatty acid composition of hyphae and spores of arbuscular mycorrhizal fungi at different growth stages. Mycological Research. 2000 Apr 1;104(4):429–34.

36. Luginbuehl LH, Menard GN, Kurup S, Erp HV, Radhakrishnan GV, Breakspear A, et al. Fatty acids in arbuscular mycorrhizal fungi are synthesized by the host plant. Science. 2017;356(6343):1175–8.

37. Sawers RJH, Svane SF, Quan C, Grønlund M, Wozniak B, Gebreselassie MN, et al. Phosphorus acquisition efficiency in arbuscular mycorrhizal maize is correlated with the abundance of root-external hyphae and the accumulation of transcripts encoding PHT1 phosphate transporters. New Phytologist. 2017 Apr 1;214(2):632–43.

38. Valentine AJ, Osborne BA, Mitchell DT. Interactions between phosphorus supply and total nutrient availability on mycorrhizal colonization, growth and photosynthesis of cucumber. Scientia Horticulturae. 2001 May 4;88(3):177–89.

39. Bakken LR, Olsen RA. Buoyant densities and dry-matter contents of microorganisms: conversion of a measured biovolume into biomass. Appl Environ Microbiol. 1983 Apr;45(4):1188–95.

40. Konvalinková T, Püschel D, Řezáčová V, Gryndlerová H, Jansa J. Carbon flow from plant to arbuscular mycorrhizal fungi is reduced under phosphorus fertilization. Plant and Soil. 2017 Oct 1;419(1):319–33.

41. Ramirez-Flores MR. Caracterización genética de mutantes de HUN e IXBA, ortólogos de maíz de los canales de potasio CASTOR y POLLUX. [Mexico]: Instituto Politecnico Nacional; 2015.

42. Warembourg FR. Estimating the true cost of dinitrogen fixation by nodulated plants in undisturbed conditions. Can J Microbiol. 1983 Aug 1;29(8):930–7.

43. Links A, Beans DE, Storage G, Know H, Fertility S, Improvement G. Making Data-Driven Decisions on Soybean Inoculation.

44. Ulzen J. Optimizing legume-rhizobia symbiosis to enhance legume grain yield in smallholder farming systems in Ghana [PhD Thesis]. 2018.

45. Schnitkey G, Paulson N, Zulauf C, Swanson K, Baltz J. Fertilizer Prices, Rates, and Costs for 2023. farmdoc daily. 2022;12(148).

46. Bonilla-Cedrez C, Chamberlin J, Hijmans RJ. Fertilizer and grain prices constrain food production in sub-Saharan Africa. Nature Food. 2021 Oct 1;2(10):766–72.

47. Reynolds HL, Hartley AE, Vogelsang KM, Bever JD, Schultz PA. Arbuscular mycorrhizal fungi do not enhance nitrogen acquisition and growth of old-field perennials under low nitrogen supply in glasshouse culture. New Phytologist. 2005 Sept 1;167(3):869–80.

48. Wang XX, Wang X, Sun Y, Cheng Y, Liu S, Chen X, et al. Arbuscular Mycorrhizal Fungi Negatively Affect Nitrogen Acquisition and Grain Yield of Maize in a N Deficient Soil. Frontiers in Microbiology [Internet]. 2018;9. Available from: https://www.frontiersin.org/journals/microbiology/articles/10.3389/fmicb.2018.00418

49. Guo K, Yang J, Yu N, Luo L, Wang E. Biological nitrogen fixation in cereal crops: Progress, strategies, and perspectives. Plant Commun. 2023 Mar 13;4(2):100499.

50. Ramana Reddy Y, Ravi D, Ramakrishna Reddy Ch, Prasad KVSV, Zaidi PH, Vinayan MT, et al. A note on the correlations between maize grain and maize stover quantitative and qualitative traits and the implications for whole maize plant optimization. Field Crops Research. 2013 Sept 1;153:63–9.

51. Tamagno S, Balboa GR, Assefa Y, Kovács P, Casteel SN, Salvagiotti F, et al. Nutrient partitioning and stoichiometry in soybean: A synthesis-analysis. Field Crops Research. 2017 Jan 1;200:18–27.

52. Ojeda-Rivera JO, Barnes AC, Ainsworth EA, Angelovici R, Basso B, Brindisi LJ, et al. Designing a nitrogen-efficient cold-tolerant maize for modern agricultural systems. The Plant Cell. 2025 July 1;37(7):koaf139.

53. Blätke MA, Bräutigam A. Evolution of C4 photosynthesis predicted by constraint-based modelling. Elife. 2019 Dec 4;8.

54. Hahn A, Vonck J, Mills DJ, Meier T, Kühlbrandt W. Structure, mechanism, and regulation of the chloroplast ATP synthase. Science. 2018 May 11;360(6389).

55. Hatch MD. C4 photosynthesis: a unique elend of modified biochemistry, anatomy and ultrastructure. Biochimica et Biophysica Acta (BBA) – Reviews on Bioenergetics. 1987 Jan 1;895(2):81–106.

56. Farquhar GD, von Caemmerer S, Berry JA. A biochemical model of photosynthetic CO2 assimilation in leaves of C3 species. Planta. 1980 June 1;149(1):78–90.

57. Chiu CH, Paszkowski U. Mechanisms and Impact of Symbiotic Phosphate Acquisition. Cold Spring Harb Perspect Biol. 2019 June 3;11(6).

58. Dreyer I, Spitz O, Kanonenberg K, Montag K, Handrich MR, Ahmad S, et al. Nutrient exchange in arbuscular mycorrhizal symbiosis from a thermodynamic point of view. New Phytologist. 2019 Apr 1;222(2):1043–53.

59. Kanehisa M. The KEGG database. In: ‘In silico’simulation of biological processes: Novartis Foundation Symposium 247. Wiley Online Library; 2002. p. 91–103.

60. Su W, Kamran M, Xie J, Meng X, Han Q, Liu T, et al. Shoot and root traits of summer maize hybrid varieties with higher grain yields and higher nitrogen use efficiency at low nitrogen application rates. PeerJ. 2019;7:e7294.

61. De Vries FWTP, Witlage JM, Kremer D. Rates of Respiration and of Increase in Structural Dry Matter in Young Wheat, Ryegrass and Maize Plants in Relation to Temperature, to Water Stress and to Their Sugar Content. Annals of Botany. 1979;44(5):595–609.

62. de Oliveira Dal’Molin CG, Quek LE, Palfreyman RW, Brumbley SM, Nielsen LK. AraGEM, a genome-scale reconstruction of the primary metabolic network in Arabidopsis. Plant Physiol. 2010 Feb;152(2):579–89.

63. Li Q, Ren Y, Fu H, Li Z, Kong F, Yuan J. Cultivar differences in carbon and nitrogen accumulation, balance, and grain yield in maize. Frontiers in Plant Science [Internet]. 2022;13. Available from: https://www.frontiersin.org/journals/plant-science/articles/10.3389/fpls.2022.992041

64. diCenzo GC, Tesi M, Pfau T, Mengoni A, Fondi M. Genome-scale metabolic reconstruction of the symbiosis between a leguminous plant and a nitrogen-fixing bacterium. Nature Communications. 2020 May 22;11(1):2574.

65. Holzhütter HG. The principle of flux minimization and its application to estimate stationary fluxes in metabolic networks. European Journal of Biochemistry. 2004 July 1;271(14):2905– 22.

66. Lewis NE, Hixson KK, Conrad TM, Lerman JA, Charusanti P, Polpitiya AD, et al. Omic data from evolved E. coli are consistent with computed optimal growth from genome-scale models. Molecular Systems Biology. 2010 Jan 1;6(1):390.

67. Ramírez-Flores MR, Perez-Limon S, Li M, Barrales-Gamez B, Albinsky D, Paszkowski U, et al. The genetic architecture of host response reveals the importance of arbuscular mycorrhizae to maize cultivation. Kliebenstein DJ, Schuman MC, editors. eLife. 2020 Nov 19;9:e61701.

68. Gutjahr C, Banba M, Croset V, An K, Miyao A, An G, et al. Arbuscular Mycorrhiza–Specific Signaling in Rice Transcends the Common Symbiosis Signaling Pathway. The Plant Cell. 2008 Nov 1;20(11):2989–3005.

69. Gräler B, Pebesma E, Heuvelink G. Spatio-Temporal Interpolation using gstat. The R Journal. 2016;8(1):204–18.

70. White CM, Finney DM, Kemanian AR, Kaye JP. Modeling the contributions of nitrogen mineralization to yield of corn. Agronomy Journal. 2021 Jan 1;113(1):490–503.

71. Giovannetti M, Mosse B. AN EVALUATION OF TECHNIQUES FOR MEASURING VESICULAR ARBUSCULAR MYCORRHIZAL INFECTION IN ROOTS. New Phytologist. 1980 Mar 1;84(3):489–500.

72. Pribyl DW. A critical review of the conventional SOC to SOM conversion factor. Geoderma. 2010;156(3–4):75–83.

73. Paul EA. The nature and dynamics of soil organic matter: Plant inputs, microbial transformations, and organic matter stabilization. Soil Biology and Biochemistry. 2016;98:109–26.

74. Panagos P, De Rosa D, Liakos L, Labouyrie M, Borrelli P, Ballabio C. Soil bulk density assessment in Europe. Agriculture, Ecosystems & Environment. 2024 Apr 15;364:108907.

75. Sullivan D, Moore A, Verhoeven E, Brewer L. Baseline soil nitrogen mineralization: measurement and interpretation. 2020 Mar.

76. Zhou Y, Li Y, Liu X, Wang K, Muhammad T. Synergistic improvement in spring maize yield and quality with micro/nanobubbles water oxygation. Scientific reports. 2019;9(1):5226.

77. Bender RR, Haegele JW, Ruffo ML, Below FE. Nutrient Uptake, Partitioning, and Remobilization in Modern, Transgenic Insect-Protected Maize Hybrids. Agronomy Journal. 2013 Jan 1;105(1):161–70.

78. Heirendt L, Arreckx S, Pfau T, Mendoza SN, Richelle A, Heinken A, et al. Creation and analysis of biochemical constraint-based models using the COBRA Toolbox v.3.0. Nature Protocols. 2019 Mar 1;14(3):639–702.

79. Ku SB, Edwards GE. Oxygen inhibition of photosynthesis[: III. Temperature dependence of quantum yield and its relation to O2/CO 2 solubility ratio. Planta. 1978 Jan;140(1):1–6.

80. Long SP, East TM, Baker NR. Chilling Damage to Photosynthesis in Young Zea mays: I. EFFECTS OF LIGHT AND TEMPERATURE VARIATION ON PHOTOSYNTHETIC CO2 ASSIMILATION. Journal of Experimental Botany. 1983 Feb 1;34(2):177–88.

81. Corn: Yield by Year, US [Internet]. United States Department of Agriculture: National Agricultural Statistics Service; 2024 [cited 2024 Oct 14]. Available from: https://www.nass.usda.gov/Charts_and_Maps/Field_Crops/cornyld.php

82. Agricultural Prices [Internet]. United States Department of Agriculture: National Agricultural Statistics Service; 2024 [cited 2024 Oct 14]. Available from: https://downloads.usda.library.cornell.edu/usda-esmis/files/c821gj76b/hd76ts06f/cc08k779s/agpr0924.pdf

83. Long-Term Impact of Fertilizer Withdrawal on Corn Yield Potential and Soil Properties. Bayer Crop Science; 2024 June.

