## Supplemental Information for "Metabolic modeling predicts synergistic growth benefits between arbuscular mycorrhizal fungi and theoretical N_2_-fixing rhizobia symbiosis in maize"

Megan L. Matthews

**This PDF file includes:**

Table S1

Figures S1 to S8

**Other supporting materials for this manuscript include the following:**

Datasets S1 to S5

**Table S1:** Values and uncertainty estimates of key plant-AMF model parameters.

| **Parameter** | **Value** | **Standard Deviation** | **Reference(s)** |
| --- | --- | --- | --- |
| AMF-to-plant biomass proportion needed for maximal uptake benefits | 0.101 | 0.028^i^ | (1,2) |
| AMF-mediated P uptake benefit | 2.16 | 0.36 | (1) |
| AMF-mediated N uptake benefit | 0.23 | 0.037^ii^ | (3) |

^i^Calculated by propagating errors of hyphal length density values reported in (1) and fresh weight / dry weight conversion and density values reported in (2).

^ii^Uncertainty estimates are not available for the values reported in (3), so we arrive at this standard deviation by assuming that the coefficient of variation of the N uptake benefit value is the same as that for the P uptake benefit value.

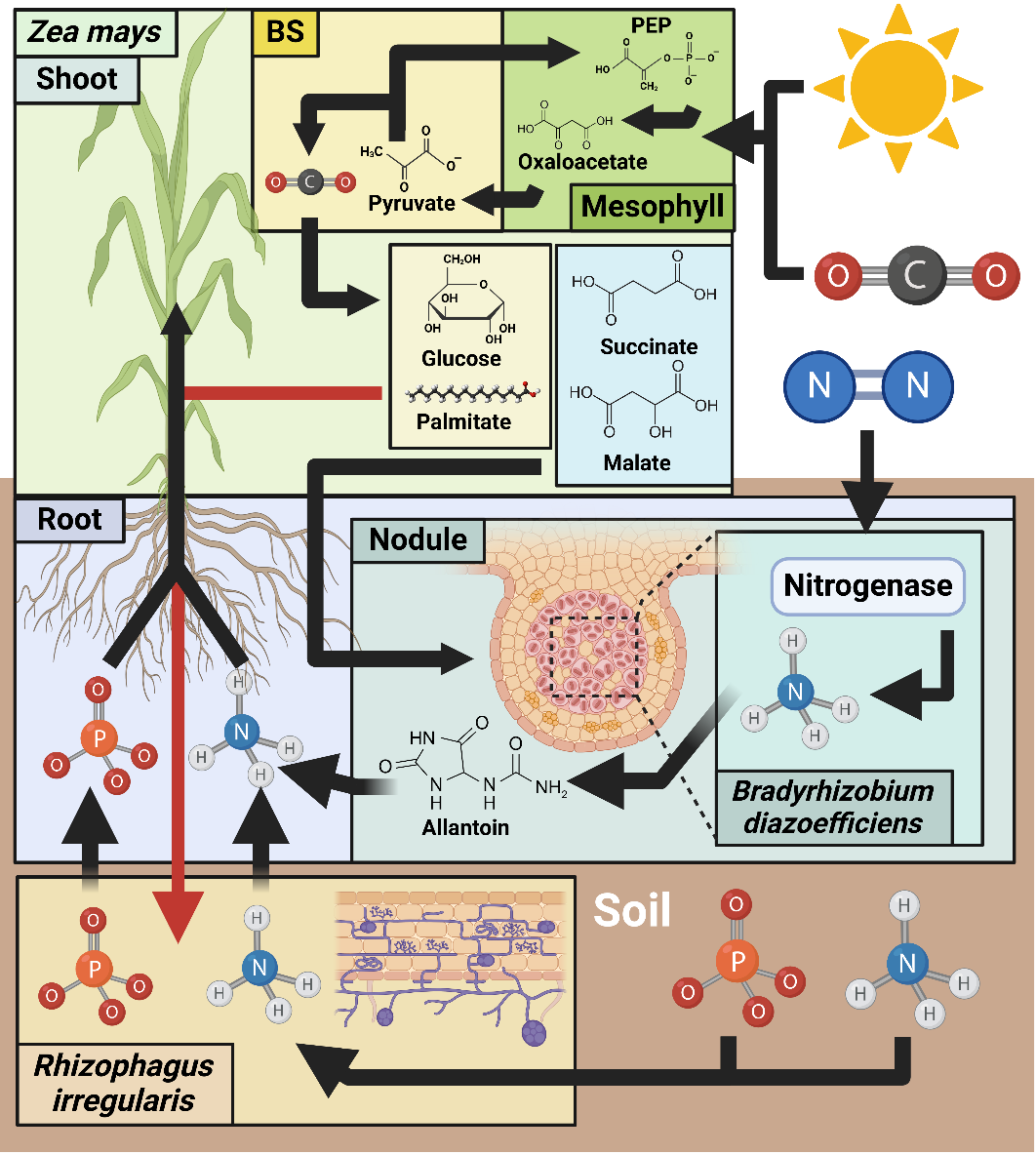

Fig. S1. Graphical representation of the major metabolic exchanges and fluxes in the three-species model developed for this study.

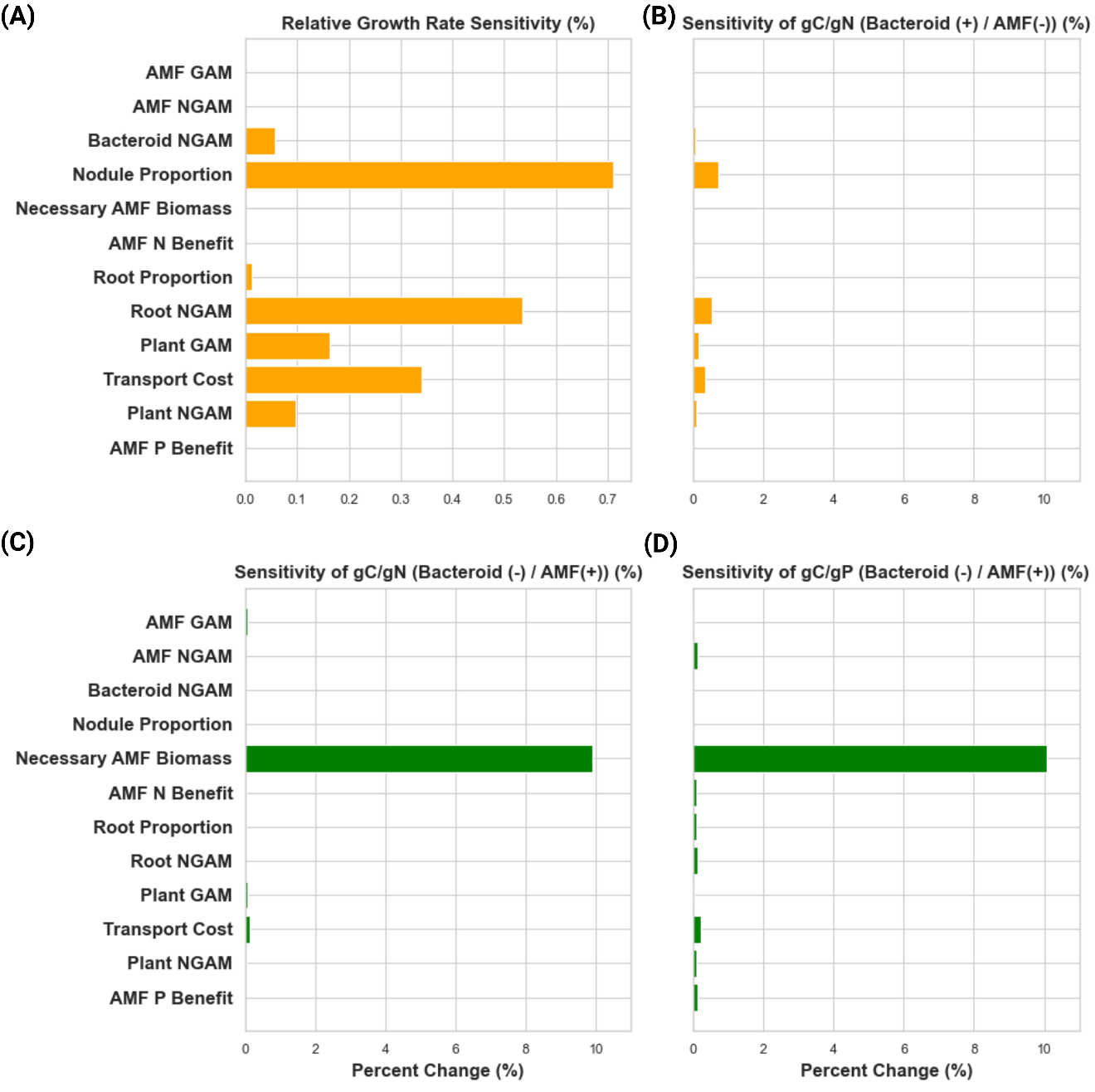

Fig. S2. Average percentage change in relative growth rate and carbon-nutrient exchange ratio resulting from a 10% increase or decrease in variable parameters across different modeling scenarios (A) Sensitivity of the relative growth rate of four models to all variable parameters. (B) Sensitivity of gC/gN estimation in the nodulated model without AMF. (C) Sensitivity of gC/gN estimation in the non-nodulated model with AMF. (D) Sensitivity of gC/gP estimation in the non-nodulated model with AMF. All gC/gN ratios are estimated with maximum P uptake and minimum N uptake, while gC/gP ratios are estimated at minimum P uptake and maximum N uptake by Zea mays.

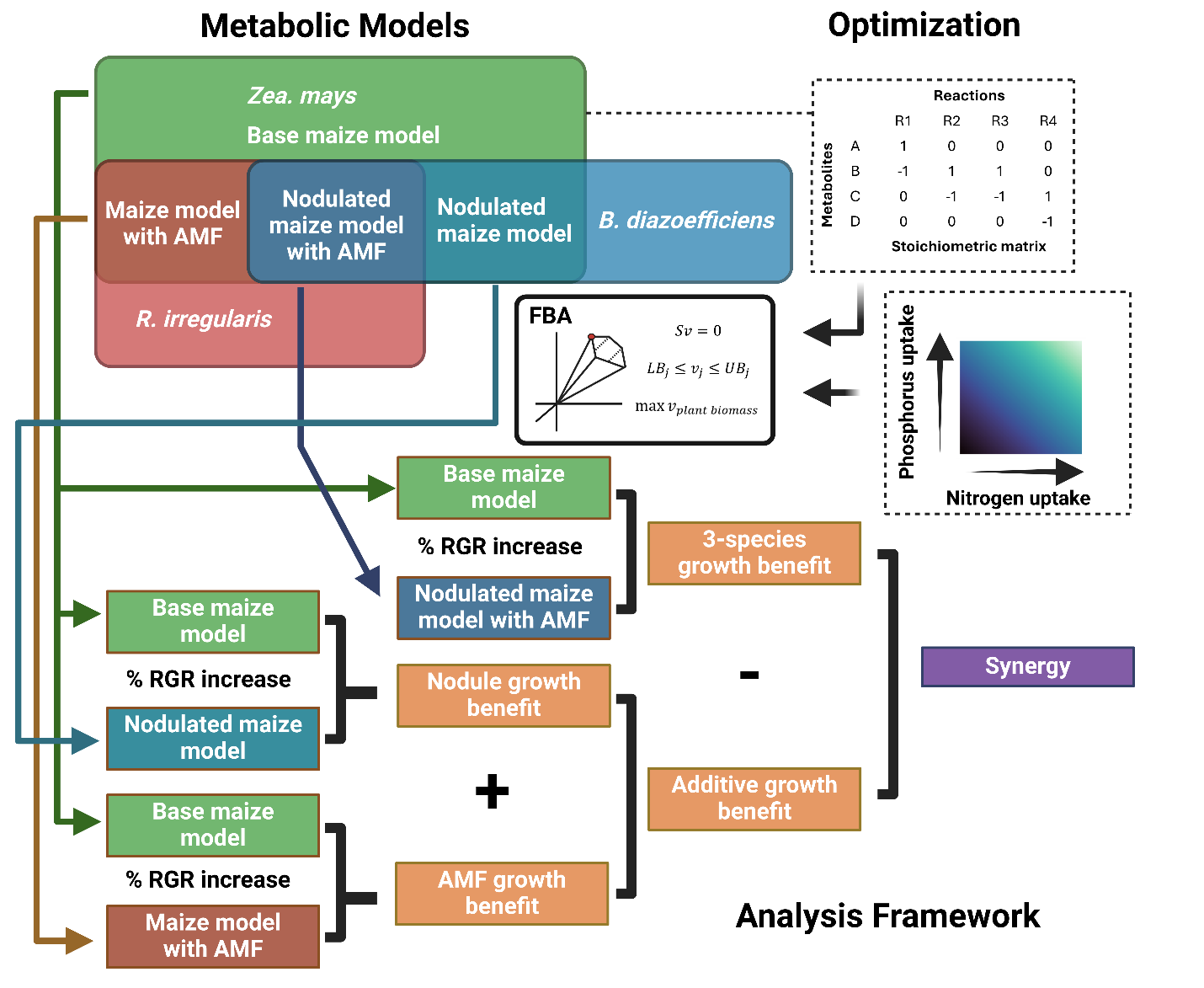

Fig. S3. Analysis framework used in this study. Through combination of a Zea mays, B. diazoefficiens, and R. irregularis model, a series of individual models representing Z. mays alone or in combination with one or two symbionts were generated. The maximized relative growth rate of Zea mays was estimated as a function of nitrogen and phosphorus uptake capacity in each of these models. Relative growth rate predictions from the base maize model and the nodulated maize model, maize model with AMF, and nodulated maize model with AMF, were compared to calculate RGR benefits and penalties under different nutrient conditions. By comparing the simple sum of the independent nodule and AMF-mediated growth benefits with the fully modeled growth benefits in the 3-species model, we calculated positive and negative synergy between the rhizobia and AMF strategies.

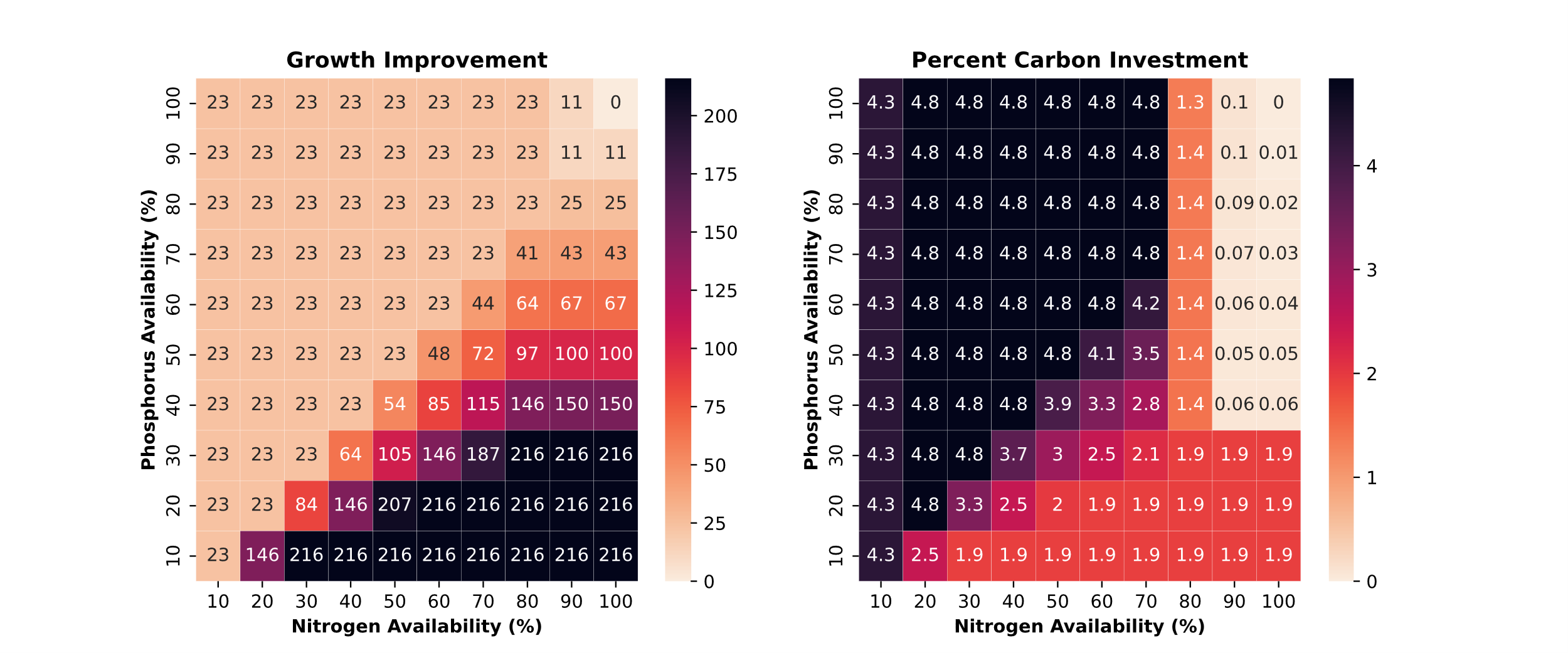

Fig. S4. Growth improvement and carbon investment predictions when comparing with- and without-AMF models when Zea mays biomass is representative of jointing C:N ratios. (A) Percentage growth improvement of with-AMF vs. without-AMF models of Zea mays as a function of phosphorus and nitrogen availability. (B) Predicted carbon investment in AMF as a percentage of net CO2 assimilation in the plant as a function of phosphorus and nitrogen availability.

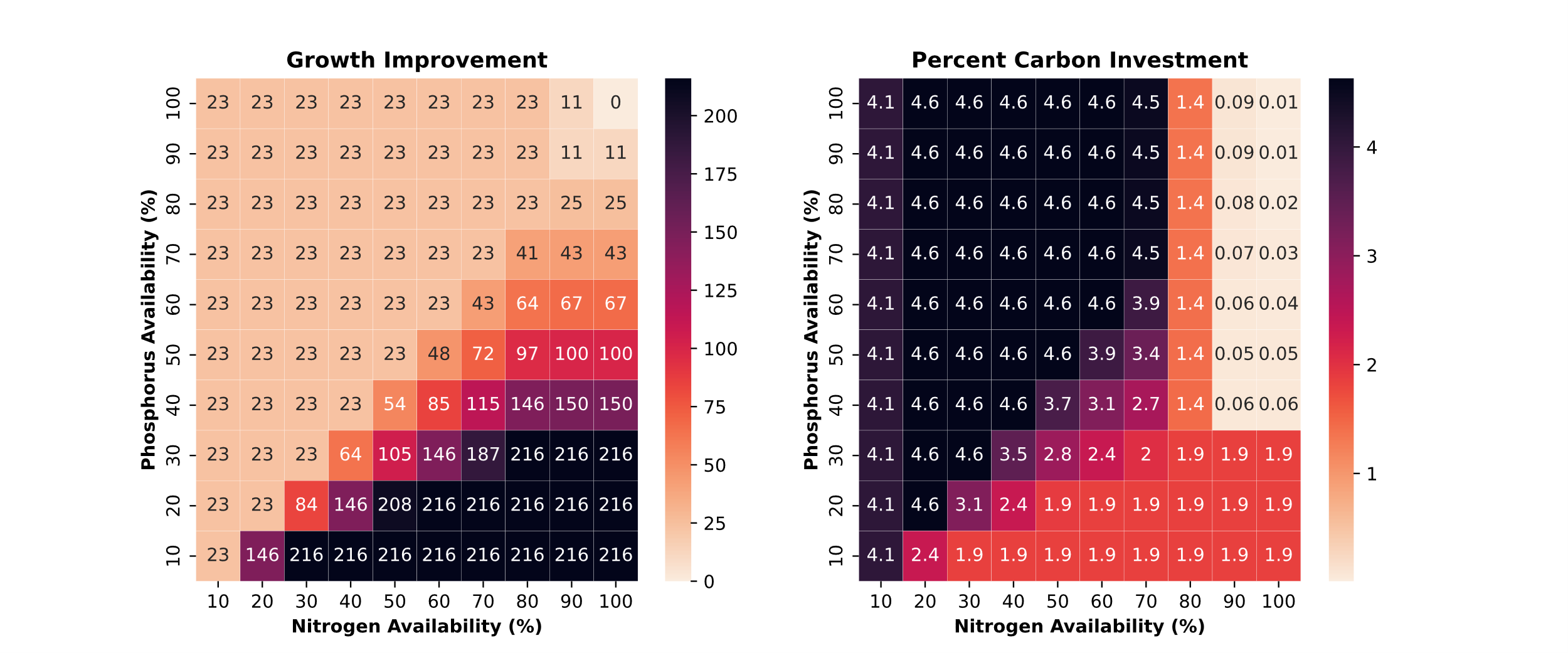

Fig. S5. Growth improvement and carbon investment predictions when comparing with- and without-AMF models when Zea mays biomass is representative of silking C:N ratios. (A) Percentage growth improvement of with-AMF vs. without-AMF models of Zea mays as a function of phosphorus and nitrogen availability. (B) Predicted carbon investment in AMF as a percentage of net CO2 assimilation in the plant as a function of phosphorus and nitrogen availability.

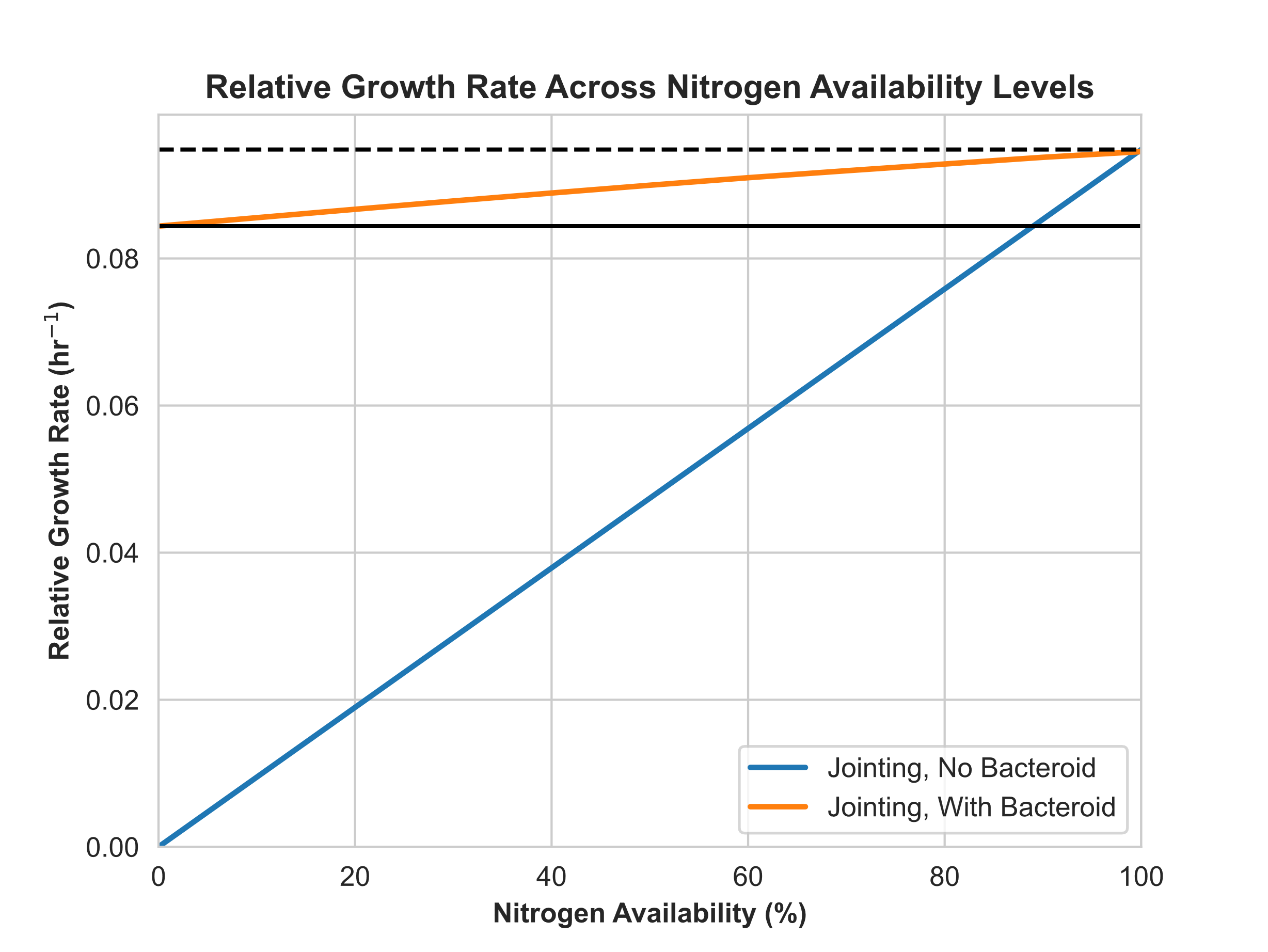

Fig. S6. Relative Growth Rate of Z. mays across a range of nitrogen availability levels for models with biomass representative of maize plants at jointing. Solid lines represent growth predictions when using the seedling biomass equation and dashed lines represent growth when using the silking biomass equation. The dashed black line represents the maximum growth rate of the non-nodulated model when it receives all of its nitrogen from the soil. The solid black line represents the maximum growth rate of the nodulated model when it fully relies on N2 fixation.

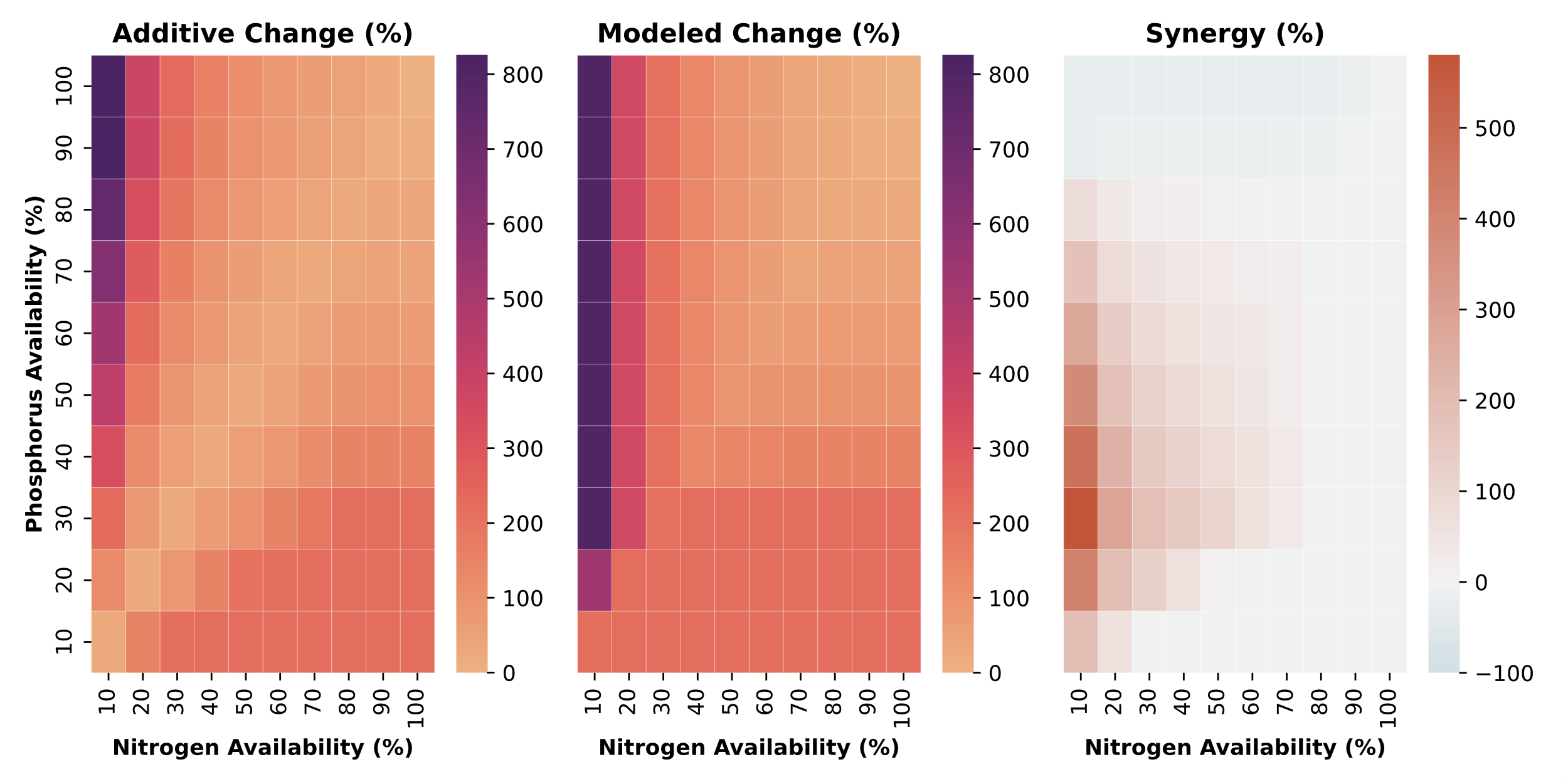

Fig. S7. Additive and modeled RGR benefits and calculated synergies and antagonisms in the Zea mays, Bradyrhizobium diazoefficiens, Rhizophagus irregularis three-species system when the Zea mays model’s biomass is representative of the C:N ratios of Zea mays plants at the jointing stage. (A) Sum of relative growth rate changes from adding B. diazoefficiens or R. irregularis to the Z. mays model under given nitrogen and phosphorus availabilities. (B) Modeled relative growth rate changes in the three-species model. (C) Synergy and antagonism between B. diazoefficiens and R. irregularis quantified as the difference between the modeled and additive growth rate changes.

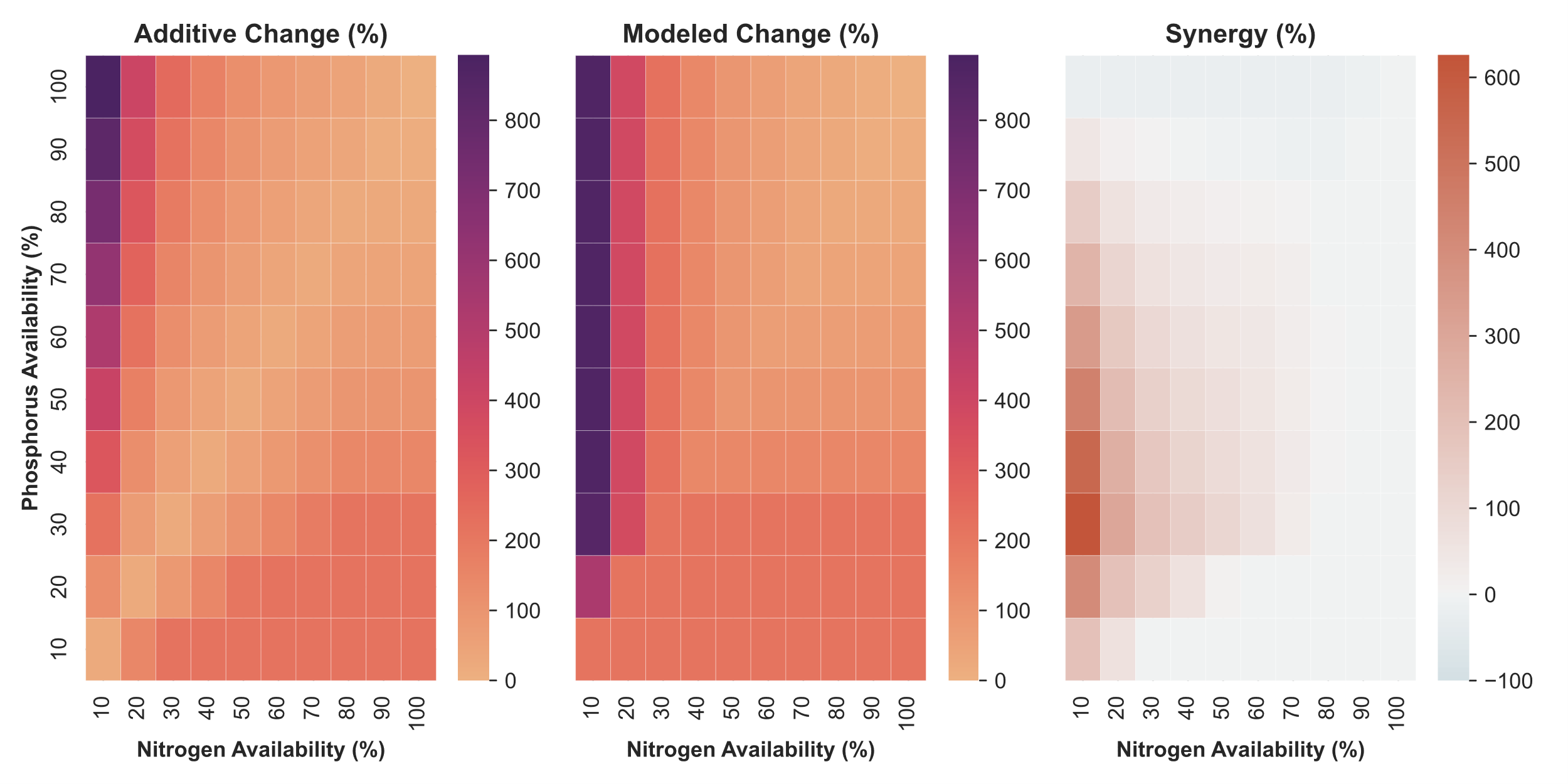

Fig. S8. Additive and modeled RGR benefits and calculated synergies and antagonisms in the Zea mays, Bradyrhizobium diazoefficiens, Rhizophagus irregularis three-species system when the Zea mays model’s biomass is representative of the C:N ratios of Zea mays plants at the silking. (A) Sum of relative growth rate changes from adding B. diazoefficiens or R. irregularis to the Z. mays model under given nitrogen and phosphorus availabilities. (B) Modeled relative growth rate changes in the three-species model. (C) Synergy and antagonism between B. diazoefficiens and R. irregularis quantified as the difference between the modeled and additive growth rate changes.

**Additional files**

Dataset S1 (separate file). List of added reactions

Dataset S2 (separate file). Efficiency and synergy results and calculations

Dataset S3 (separate file). AMF+/- field data and calculations

Dataset S4 (separate file). GAM and NGAM calculation details

Dataset S5 (separate file). Transport reaction details

Supplemental References

1.	Sawers RJH, Svane SF, Quan C, Grønlund M, Wozniak B, Gebreselassie MN, et al. Phosphorus acquisition efficiency in arbuscular mycorrhizal maize is correlated with the abundance of root-external hyphae and the accumulation of transcripts encoding PHT1 phosphate transporters. New Phytologist. 2017 Apr 1;214(2):632–43.

2.	Bakken LR, Olsen RA. Buoyant densities and dry-matter contents of microorganisms: conversion of a measured biovolume into biomass. Appl Environ Microbiol. 1983 Apr;45(4):1188–95.

3.	Valentine AJ, Osborne BA, Mitchell DT. Interactions between phosphorus supply and total nutrient availability on mycorrhizal colonization, growth and photosynthesis of cucumber. Scientia Horticulturae. 2001 May 4;88(3):177–89.
